# Ca^2+^-activated K^+^ channels reduce network excitability, improving adaptability and energetics for transmitting and perceiving sensory information

**DOI:** 10.1101/476861

**Authors:** Xiaofeng Li, Ahmad Abou Tayoun, Zhuoyi Song, An Dau, Diana Rien, David Jaciuch, Sidhartha Dongre, Florence Blanchard, Anton Nikolaev, Lei Zheng, Murali K. Bollepalli, Brian Chu, Roger C. Hardie, Patrick J. Dolph, Mikko Juusola

**Affiliations:** State Key Laboratory of Cognitive Neuroscience and Learning, Beijing Normal University, Beijing 100875, China; Department of Biomedical Science, University of Sheffield, Sheffield S10 2TN, UK; Department of Biology, Dartmouth College, Hanover, New Hampshire, NH 03755, USA; Institute of Science and Technology for Brain-Inspired Intelligence and Key Laboratory of Computational Neuroscience and Brain-Inspired Intelligence, Ministry of Education, Fudan University, Shanghai 200433, China; Department of Physiology, Development, and Neuroscience, University of Cambridge, Cambridge CB2 3DY, UK

## Abstract

Ca^2+^-activated K^+^ channels (BK and SK) are ubiquitous in synaptic circuits, but their role in network adaptation and sensory perception remains largely unknown. Using electrophysiological and behavioral assays and biophysical modelling, we discover how visual information transfer in mutants lacking the BK channel (*dSlo*^−^), SK channel (*dSK*^−^) or both (*dSK*^−^;;*dSlo*^−^) is shaped in the female fruit fly (*Drosophila melanogaster*) R1-R6 photoreceptor-LMC circuits (R-LMC-R system) through synaptic feedforward-feedback interactions and reduced R1-R6 *Shaker* and *Shab* K^+^ conductances. This homeostatic compensation is specific for each mutant, leading to distinctive adaptive dynamics. We show how these dynamics inescapably increase the energy cost of information and promote the mutants’ distorted motion perception, determining the true price and limits of chronic homeostatic compensation in an *in vivo* genetic animal model. These results reveal why Ca^2+^-activated K^+^ channels reduce network excitability (energetics), improving neural adaptability for transmitting and perceiving sensory information.

**Significance statement:** In this study, we directly link *in vivo* and *ex vivo* experiments with detailed stochastically operating biophysical models to extract new mechanistic knowledge of how *Drosophila* photoreceptor-interneuron-photoreceptor (R-LMC-R) circuitry homeostatically retains its information sampling and transmission capacity against chronic perturbations in its ion-channel composition, and what is the cost of this compensation and its impact on optomotor behavior. We anticipate that this novel approach will provide a useful template to other model organisms and computational neuroscience, in general, in dissecting fundamental mechanisms of homeostatic compensation and deepening our understanding of how biological neural networks work.

## Introduction

Ca^2+^-activated K^+^ channels are widely expressed in both the visual system and CNS and play important roles in cell physiology, such as modulating neuronal excitability and neurotransmitter release. Based upon their kinetics, pharmacological and biophysical properties, these channels can be divided into two main types: the “small”-(SK; 2-20 pS) and “big”-conductance (BK; 200-400 pS) channels. The SK channels are solely Ca^2+^-activated (Sah, 1996; Faber and Sah, 2003; Stocker, 2004; Salkoff, 2006), while BK channels are both Ca^2+^- and voltage-dependent. At synapses, SK channels form negative feedback loops with Ca^2+^ sources and are therefore essential regulators of synaptic transmission (Faber et al., 2005; Ngo-Anh et al., 2005). The functional role of BK channels in synaptic activities is less well understood, with various effects of blocking BK channels on neurotransmitter release having been reported (Fettiplace and Fuchs, 1999; Ramanathan et al., 1999; Xu and Slaughter, 2005).

Although Ca^2+^-activated K^+^ channels – through regulation of synaptic transmission between retinal neurons – seem to have conserved roles in early vertebrate (Shatz, 1990; Wang et al., 1999; Klocker et al., 2001; Pelucchi et al., 2008; Clark et al., 2009; Grimes et al., 2009) and invertebrate vision (Abou Tayoun et al., 2011), it has been difficult to work out how these channels advance *in vivo* circuit functions and what are their evolutionary benefits. This is because homeostatic processes that regulate electrical activity in neurons, in part, make communication in circuits surprisingly fault-tolerant against perturbations (Lemasson et al., 1993; Marder and Goaillard, 2006). Thus, the physical consequences of altering K^+^ channel densities and those of homeostatic compensation are interconnected. Because *Drosophila* has single *SK* (*dSK*) and *BK* (*dSlo*) genes, electrophysiologically accessible photoreceptors and interneurons (large monopolar cells, LMCs) (Juusola and Hardie, 2001a; Zheng et al., 2006) with stereotypical connectivity (Meinertzhagen and O’Neil, 1991; Rivera-Alba et al., 2011), and readily quantifiable optomotor behavior (Blondeau and Heisenberg, 1982; Juusola et al., 2017), it provides an excellent model system to characterize how Ca^2+^-activated K^+^ channels affect circuit functions and the capacity to see. Importantly, *Drosophila* photoreceptors and LMCs express both *dSK* and *dSlo* genes (Abou Tayoun et al., 2011; Davis et al., 2018). Here, we study to what extent intrinsic perturbations of missing one or both of these K^+^ channels, through gene-deletion, can be neutralized by homeostatic processes trying to sustain normal network functions, and what is the price of this compensation.

By using electrophysiological and behavioral assays and biophysical modelling, we uncover why Ca^2+^-activated K^+^ channels improve communication between photoreceptors and Large Monopolar Cells (LMCs), which in the fly eye lamina network form stereotypical columns of feedforward and feedback synapses (R-LMC-R system) that process and route visual information to the fly brain. We show that although the loss of SK and BK channels does not diminish *Drosophila* photoreceptors’ information sampling capacity *in vivo*, it homeostatically reduces other K^+^ currents and overloads synaptic-feedback from the lamina network. This makes communication between the mutant photoreceptors and LMCs inefficient, consuming more energy and distorting visual information flow to the brain. Thus, homeostatic compensation of missing SK and BK channels within the lamina network is suboptimal and comes with an unavoidable cost of reduced adaptability and altered (accelerated or decelerated) vision, which thereby must contribute to the mutant flies’ uniquely tuned optomotor behaviors.

These results quantify the benefits of Ca^2+^-activated K^+^ channels in improving robustness, economics and adaptability of neural communication and perception.

## Materials and Methods

### *Drosophila melanogaster* strains and rearing

*w*+; +; *dSlo4* (Gift from Nigel Atkinson laboratory, identifiers: CG10693, FBgn0003429).

*w*+; +; *dSlo4/dSlo18* (Gift from Allen Shearn laboratory, identifiers: Dmel\ash218, FBal0057820).

*w*+; *dSK*-; + (Patrick Dolph laboratory, identifiers: CG10706, FBgn0029761).

*w*+; *dSK*-; +; *dSlo4* (in house).

*w*+*dSK*-; +; *dSlo4/dSlo18* (in house).

The flies were maintained in the stock as:

*w*+; +; *Slo4/TM6*

*w*+ *dSK*-; +; +

*w*+*dSK*-; +; *dSlo4/TM6*

*w*+; +; *dSlo18/TM6*

The *dSK*^−^ and *UAS-dSKDN* alleles were prepared as described earlier (Abou Tayoun et al., 2011). *Df7753* or *Df(1)Exel6290* line was obtained from Bloomington *Drosophila* stock center.

*dSlo^4^* null allele (Atkinson et al., 1991) was kindly provided by Dr. Nigel Atkinson. *dSlo^4^* mutants appear often unhealthy, with the dSlo channel being expressed both in muscles and the brain (due to its 2 independent control regions), making them hesitant fliers (Atkinson et al., 2000). Therefore, we generated transheterozygotes *dSlo^4^/dSlo^18^*, facilitating the flight simulator experiments. *dSlo^4^* and *dSlo^18^* (also called *ash2^18^*) are both mutations of slowpoke (Lajeunesse and Shearn, 1995; Atkinson et al., 2000). But slowpoke has multiple promoters: *dSlo^4^* is a loss of function, whereas *dSlo^18^* affects promoter C0 and C1 (neural-specific) yet leaves C2 promoter intact. *dSlo^18^* produces a functional channel in the muscle, thereby mostly rescuing the flight deficits. *dSlo^18^* only affects the brain control region and is homozygous lethal, and thus, both *dSlo^4^* and *dSlo^18^* were maintained over a TM6b balancer. For experimental flies, *dSlo^4^/TM6* or *dSK*;;*dSlo^4^/TM6* were crossed to *dSlo^18^* and we selected against the *TM6* balancer. When combined in a *dSlo^4^/dSlo^18^*, the mutations only affect the expression of *dSlo* in the brain. All the flies were previously outcrossed to a common Canton-S background, which was the wild-type control. The overall yield of *dSlo*^−^ mutants was lower than for the other flies, with the surviving adults flies being typically smaller, which suggested that homozygotic *dSlo*^−^ mutants were less healthy.

*Drosophila* were raised on molasses based food at 18 °C, under 12:12 h light:dark conditions. Prior to the experiments, the flies were moved to the laboratory (~21 °C) overnight or kept in a separate incubator at 25 °C. All electrophysiology (intracellular, electroretinogram and whole-cell recordings) was conducted at 20 ± 1 °C and optomotor behavior experiments at 21 ± 1 °C. During *in vivo* recordings, the fly temperature was feedback-controlled by a Peltier-system (Juusola and Hardie, 2001a; Juusola et al., 2016). Moreover, the theoretical model simulations of the R-LMC-R system (see below) were also calculated for 20 °C, by adjusting the Q_10_ of phototransduction reactions and membrane properties accordingly (Juusola and Hardie, 2001a; Song et al., 2012). Thus, by retaining effectively the same temperature for experiments and theory, we could compare directly the wild-type and mutant electrophysiology to their respective model predictions and optomotor behaviors.

Because the intracellular response dynamics of *dSlo^4^* and *dSlo^4^/dSlo^18^* R1-R6 photoreceptors and LMCs, respectively, appeared consistently similar, differing in the same way from the wild-type responses, these responses were pooled in the main results (Figs 3–12). For the same reason, the corresponding responses of *dSK*^−^;;*dSlo^4^* and *dSK*^−^;;*dSlo^4^/dSlo^18^* R1-R6 and LMCs were also pooled.

#### Why dSK and dSlo expression in photoreceptors or LMCs was not manipulated using Gal4-drivers

When using mutant animals, it is standard practice to use cell-specific transgenic rescues to show that the described phenotype is causally linked to the used mutations (rather than genetic background effects linked to the mutation), and/or to complement classical mutants with cell-specific RNAi knockdown. This is normally done by using cell-specific Gal4 drivers to control expression of transgenes or RNAis. However, for the current study, we deemed these methods unviable. For LMCs, we could not use them because for technical reasons our recordings mix L1 and L2 cell types; see ***In vivo intracellular recordings*** below. Conversely, for common Gal4 photoreceptor lines (e.g. GMR, longGMR), our tests indicate that these compromise development (Bollepalli et al., 2017), causing reductions in sensitivity, dark noise, potassium currents, and cell size and capacitance, as well as extreme variations in sensitivity between cells (Fig. 1). We have also found that another commonly used line (Rh1-Gal4), although not causing the same developmental abnormalities, leads to highly variable (~100-fold) UAS-transgene expression levels from cell to cell (Fig. 1F). In our hands, we have also found that effective RNAi in the photoreceptors is only reliably achieved with the strong GMR or longGMRGal4 drivers with their attendant developmental abnormalities (Fig. 1). Thus, the use of Gal4 drivers would be expected to induce experimental variability, altering photoreceptor output dynamics far more than the contribution of missing dSlo- or dSK-channels. Such controls would therefore increase uncertainty rather than reduce it, making their use here scientifically unsound.

**Figure 1.**
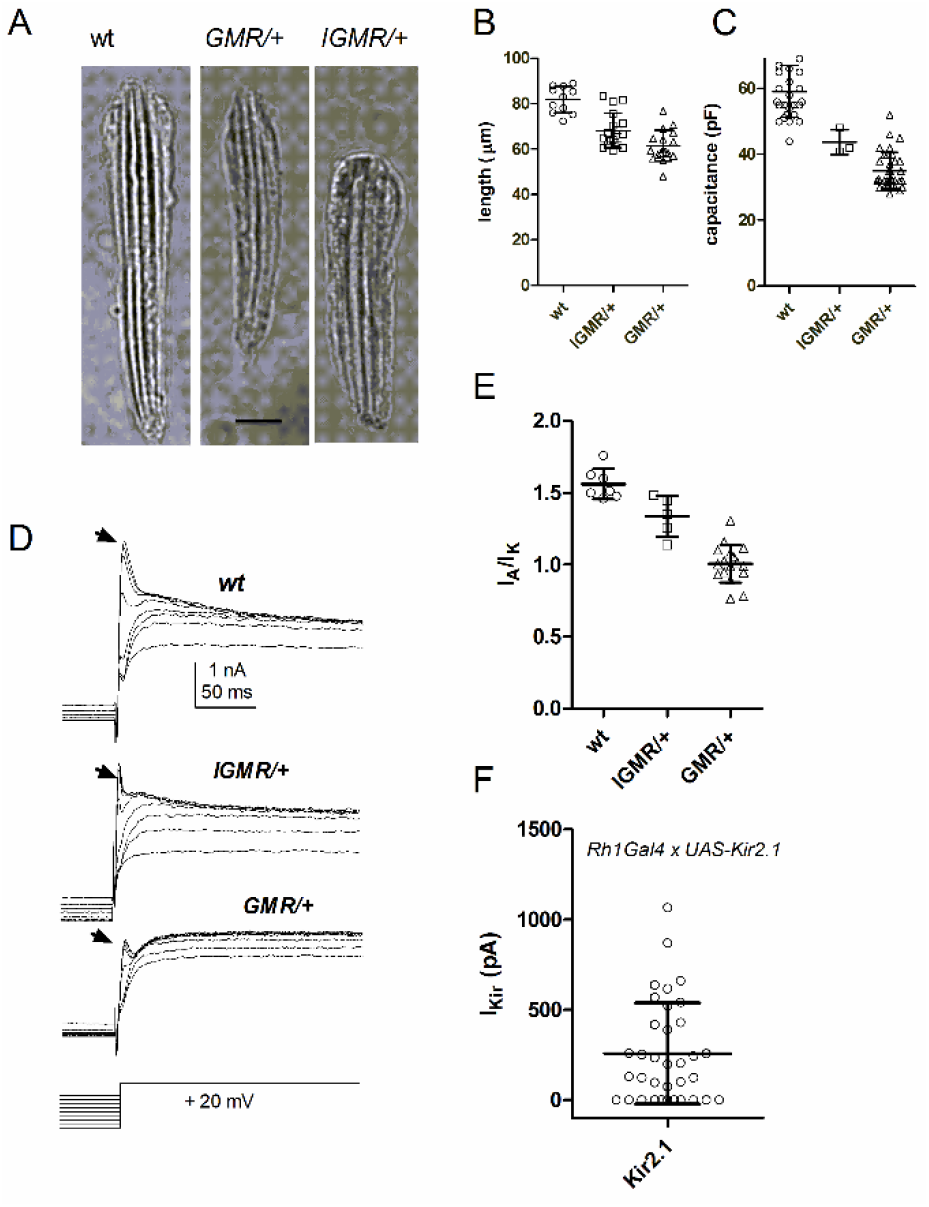
Gal4-controls alter photoreceptor structure and function. ***A***, Images of dissociated ommatidia from wild-type (wt) and flies expressing one copy of GMRGal4 (GMR/^+^) and longGMRGal4 (IGMR/^+^). Scale bar 10 μm. ***B***, Ommatidial length (wt, n=11; longGMR, n=18; GMR/+, n=17; p<0.0001) and *C*, whole-cell capacitance (wt, n=30; longGMR, n=3 [p=0.002]; GMR, n=35 [p<0.0001]), ***C***, are substantially reduced by expression of one copy of both GMRGal4 and longGMRGal4. ***D***, Native *Shaker* (I_A_ arrows) and delayed rectifier, *Shab* (I_Ks_) current profiles are altered in both GMRGal4 and longGMRGal4 flies. ***E***, Summary data: ratio of *Shaker* (I_A_) to *Shab* (I_Ks_) currents (wt, n=7; longGMR/+, n = 5 [p=0.01]; GMR/, n=17 [p=0.002]). ***F***, Expression levels of a K channel transgene (UAS-Kir2.1) driven by Rh1Gal4 was extremely variable (inward rectifier currents measured from n = 36 cells), with ~30% of photoreceptors cells showing no detectable expression at all. ***B-C*** and ***E-F***: Mean ± SD, two-tailed t-test. All significant at p<0.05 also on ANOVA plus Tukey’s Multiple comparison.

### Electrophysiology and Analysis

#### Electroretinograms (ERGs)

ERGs were recorded from intact flies following the standard procedures (Dau et al., 2016). ≤1 week old adult female *Drosophila* were fixed into a conical holder (Juusola and Hardie, 2001b; Juusola et al., 2016), using low melting point beeswax, and stimulated by 1 s light pulses from a green (560 nm) LED with the brightest effective intensity, estimated to be ~5 × 10^6^ effective photons/photoreceptor/s. Both recording and reference electrodes were filled with *Drosophila* ringer (in mM): 120 NaCl, 5 KCl, 1.5 CaCl_2_, 4 MgCl_2_, 20 proline, and 5 alanine. The recording electrode was positioned to touch the cornea and the indifferent electrode the head capsule near the ocelli. Recorded signals were low-pass filtered at 200 Hz and amplified via a npi SEC-10LX amplifier (npi Electronics, Germany).

A wild-type ERG comprises two main components: a slow component and transients coinciding with changes in light stimuli (Heisenberg, 1971). The slow component (or maintained background potential) is attributed to photoreceptor output and has the inverse waveform of photoreceptors’ intracellular voltage responses, while on- and off-transients originate from the postsynaptic cells in the lamina (Coombe, 1986). We further plotted the ERGs as dynamic spectra (Fig. 8*H*) to highlight how their oscillation frequencies changed in respect to light stimulation (Wolfram and Juusola, 2004).

#### Whole-Cell Recordings

Dissociated ommatidia were prepared from recently eclosed adult female flies and transferred to a recording chamber on an inverted Nikon Diaphot microscope (Hardie et al., 2002). The control bath solution contained 120 mM NaCl, 5 mMKCl, 10mM N-Tris-(hydroxymethyl)-methyl-2-amino-ethanesulphonic acid (TES), 4 mM MgCl_2_, 1.5 mM CaCl_2_, 25 mMproline, and 5 mM alanine. Osmolarity was adjusted to ~283 mOsm. The intracellular solution used in the recording pipette was composed of 140 mM K^+^ gluconate, 10 mM TES, 4 mM Mg^2+^ ATP, 2 mM MgCl_2_, 1 mM NAD, and 0.4 mM Na^+^ GTP. Data were recorded with Axopatch 1-D or 200 amplifiers and analyzed with pClamp software (Axon Instruments). Cells were stimulated by a green-light-emitting diode with intensities calibrated in terms of effectively absorbed photons by counting quantum bumps at low intensities in wild-type flies.

#### In vivo intracellular recordings

3-7 days old (adult) female flies were used in the experiments; the female flies are larger than the males, making the recordings somewhat easier. A fly was fixed in a conical fly-holder with beeswax, and a small hole (6-10 ommatidia) for the recording microelectrode entrance was cut in its dorsal cornea and Vaseline-sealed to protect the eye (Juusola and Hardie, 2001b; Zheng et al., 2006). Sharp quartz and borosilicate microelectrodes (Sutter Instruments), having 120–200 MΩ resistance, were used for intracellular recordings from R1-R6 photoreceptors and large monopolar cells (LMCs). These recordings were performed separately; with the electrodes filled either with 3 M KCl solution for photoreceptor or 3 M potassium acetate with 0.5 mM KCl for LMC recordings (Juusola et al., 1995b; Zheng et al., 2006), to maintain chloride battery. A reference electrode, filled with fly ringer, was gently pushed through ocelli ~100 μm into the head, in which temperature was kept at 19 ± 1°C by a Peltier device (Juusola and Hardie, 2001a).

Only stable high-quality recordings were included. In darkness, R1-R6s’ maximum responses to saturating bright pulses were characteristically >40 mV (wild-type, all mutants); the corresponding LMC recordings showed resting potentials <-30 mV and 10-40 mV maximum response amplitudes (wild-type and all mutants). Although the large maximum response variation is typical for *Drosophila* intracellular LMC recordings, their normalized waveforms characteristically display similar time-courses and dynamics (Nikolaev et al., 2009; Zheng et al., 2009). The smaller and more frequent responses are likely from LMC somata. These have larger diameters than the small and narrow LMC dendrites, in which responses should be the largest but the hardest to record from (Nikolaev et al., 2009; Zheng et al., 2009; Wardill et al., 2012). LMC subtypes were not identified, but most recordings were likely from L1 and L2 as these occupy the largest volume. Occasionally, we may have also recorded from other neurons or glia, which receive histaminergic inputs from photoreceptors (Shaw, 1984; Zheng et al., 2006; Zheng et al., 2009; Rivera-Alba et al., 2011). But because the selected recordings shared similar hyperpolarizing characteristics, LMC data for each genotype were analyzed together. Such pooling is further justified by the Janelia Farm gene expression data (Davis et al., 2018), which shows that both dSK and dSlo genes are rather highly expressed (in transcripts/million units) across all LMC types, and that all of these cells are expressing both genes.

Light stimulation was delivered to the studied cells at center of its receptive field with a high-intensity green LED (Marl Optosource, with peak emission at 525 nm), through a fiber optic bundle, fixed on a rotatable Cardan arm, subtending 5° as seen by the fly. Its intensity was set by neutral density filters (Kodak Wratten) (Juusola and Hardie, 2001b); the results are shown for dim (estimated to be ~600), medium (~6 × 10^4^) and bright luminance (~6 × 10^5^ photons/s); or log −3, log −1 and log 0, respectively.

Voltage responses were amplified in current-clamp mode using 15 kHz switching rate (SEC-10L single-electrode amplifier; NPI Electronic, Germany). The stimuli and responses were low-pass filtered at 500 Hz (KemoVBF8), and sampled at 1 or 10 kHz. The data were re-sampled/processed off-line at 1-2 kHz for the analysis. Stimulus generation and data acquisition were performed by custom-written Matlab (MathWorks, Natick, MA) programs: BIOSYST (Juusola and Hardie, 2001b; Juusola and de Polavieja, 2003).

#### Data Analysis

The signal was the average of consecutive 1,000 ms long voltage responses to a repeated light intensity time series, selected from the naturalistic stimulus (NS) library (van Hateren, 1997), and its power spectrum was calculated using Matlab’s Fast Fourier Transform (FFT) algorithm. First 10-20 responses were omitted because of their adaptive trends, and only approximately steady-state adapted responses were analyzed. The noise was the difference between individual responses and the signal, and its power spectra were calculated from the corresponding traces (Juusola et al., 1994). Thus, n trials (with n = 20), gave one signal trace and n noise traces. Both signal and noise data were chunked into 50% overlapping stretches and windowed with a Blackman-Harris-term window, each giving three 500-point-long samples. This gave 60 spectral samples for the noise and three spectral samples for the signal, which were averaged, respectively, to improve the estimates. *SNR*(*f*), of the recording or simulation was calculated from their signal and noise power spectra, <|*S*(*f*)|^2^> and <|*N*(*f*)|^2^>, respectively, as their ratio, where || denotes the norm and <> the average over the different stretches (Juusola and Hardie, 2001b; Juusola and de Polavieja, 2003; Song and Juusola, 2014).

Information transfer rates, *R*, for each recording were estimated by using the Shannon formula (Shannon, 1948), which has been shown to obtain robust estimates for these types of continuous signals (Juusola and de Polavieja, 2003; Song and Juusola, 2014; Juusola et al., 2017). We analyzed steady-state-adapted recordings and simulations, in which each response (or stimulus trace) is expected to be equally representative of the underlying encoding (or statistical) process. From *SNR*(*f*), the information transfer rate estimates were calculated as follows:

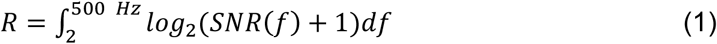

with the integral upper and lower bounds resulting from 1 kHz sampling rate and 500 points window size, respectively. The underlying assumptions of this method and how the number and resolution of spectral signal and noise estimates and the finite size of the used data can affect the resulting information transfer rate estimates have been analyzed before (van Hateren, 1992b; Juusola and de Polavieja, 2003; Song and Juusola, 2014) and are further discussed in (Juusola et al., 2017).

Using some longer recording series (to 50 stimulus repetitions), we further tested these *R* estimates against those obtained by the triple extrapolation method (Juusola and de Polavieja, 2003). This method, unlike SNR analysis, requires no assumptions about the signal and noise distributions or their additivity. Voltage responses were digitized by sectioning them into time intervals, *T*, that were subdivided into smaller intervals *t* = 1 ms. In the final step, the estimates for the entropy rate, *R_S_*, and noise entropy rate, *R_N_*, were then extrapolated from the values of the experimentally obtained entropies to their successive limits, as in (Juusola and de Polavieja, 2003):

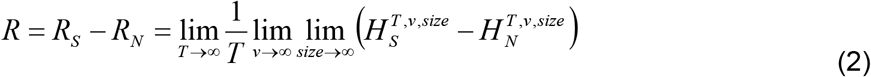

where *T* is the length of the ‘words’, *ν* the number of voltage levels (in digitized amplitude resolution) and the *size* of the data file. The difference between the entropy and noise entropy rates is the rate of information transfer, *R* (Shannon, 1948; Juusola and de Polavieja, 2003). Again, as shown before for comparable data (Song and Juusola, 2014; Dau et al., 2016; Juusola et al., 2017), both methods gave similar *R* estimates, implying that the Shannon method (Eq. 1) estimates were unbiased.

As expected, information transfer rates at 20 °C were lower than those at 25 °C (Song and Juusola, 2014; Juusola et al., 2017), which is *Drosophila’s* preferred temperature (Sayeed and Benzer, 1996). Presumably, because of the tightly-compartmentalized enzymatic reactions inside each of its 30,000 microvilli (phototransduction/photon sampling units), the Q_10_ of a *Drosophila* R1-R6’s information transfer is high for many light stimuli; ≥4 for bright 200 Hz Gaussian white-noise stimulation (Juusola and Hardie, 2001a). Whereas, the Q_10_ of simple diffusion-limited reactions, such as ion channel currents, is lower, ~2 (Lamb, 1984; Juusola and Hardie, 2001a). Critically here, stochastic R1-R6 model simulations imply that warming accelerates microvilli recovery from their previous light-activation by shortening their refractory period (Song and Juusola, 2014). Therefore, for many bright fast-changing light patterns, a warm R1-R6 transduces characteristically more photons to quantum bumps than a cold one. And, with more bumps summing up bigger and faster macroscopic responses, extending their reliability to higher stimulus frequencies, information transfer increases (Juusola and Hardie, 2001a; Juusola et al., 2016; Juusola and Song, 2017).

### Behavioral Experiments and Analysis

In the flight simulator experiments, we used 3-7 days old female flies, reared in 12:12 h dark:light cycle. A flying fly, tethered from the classic torque-meter (Tang and Guo, 2001), which fixed its head in a rigid position and orientation, was lowered by a manipulator in the center of a black-white cylinder (spectral full-width: 380-900 nm). It saw a continuous (360°) stripe-scene. After viewing the still scene for 1 s, it was spun to the counter-clockwise by a linear stepping motor for 2 s, stopped for 2 s, before rotating to clock-wise for 2 s, and stopped again for 1 s. This 8 s stimulus was repeated 10 times and each trial, together with the fly’s yaw torque responses, was sampled at 1 kHz and stored for later analysis (Wardill et al., 2012). Flies followed the scene rotations, generating yaw torque responses (optomotor responses to right and left), the strength of which presumably reflects the strength of their motion perception (Götz, 1964). The moving stripe scenes had: azimuth ±360°; elevation ±45°; wavelength 14.4° (coarse) and 3.9° (fine-grained = hyperacute); contrast 1.0, as seen by the fly. The scene was rotated at 45°/s (slow) of 300°/s (fast).

### Biophysical Models for Estimating Wild-type and Mutant R1-R6s’ Energy Consumption

Our published (Song et al., 2012) and extensively tested (Song and Juusola, 2014; Juusola et al., 2015; Juusola et al., 2017; Song and Juusola, 2017) *Drosophila* R1-R6 photoreceptor model (Figs *2A-D*), was used to simulate both the wild-type and mutant voltage responses to naturalistic light intensity time series. It has four modules: (1) random photon absorption model, which regulates photon absorptions in each microvillus, following Poisson statistics (Fig. *2C*, green); (2) stochastic quantum bump (QB) model, in which stochastic biochemical reactions inside a microvillus captures and transduces the energy of photons to variable QBs or failures (Figs 2*A-B*); (3) summation model, in which QBs from 30,000 microvilli integrate the macroscopic light-induced current (LIC) response (Fig. 2*C*, blue); and (4) Hodgkin–Huxley (HH) model of the photoreceptor plasma-membrane (Niven et al., 2003; Vähäsöyrinki et al., 2006; Song et al., 2012), which transduces LIC into voltage response (Fig. 2*D*). The model’s open-source Matlab code can be downloaded from GitHub (the links below).

**Figure 2.**
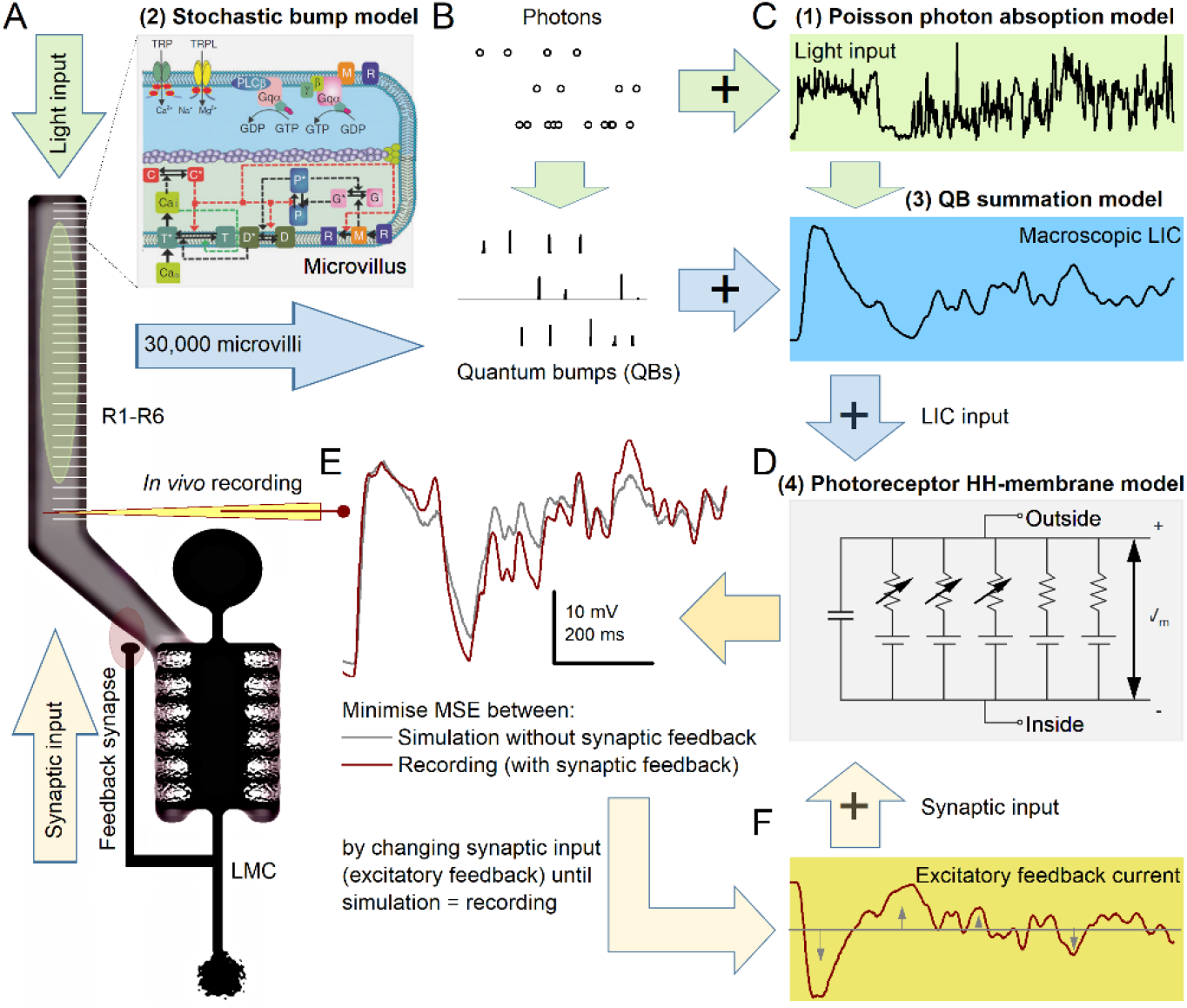
Schematic R1-R6 model structure with excitatory synaptic feedback from lamina interneurons (LMCs: L2 and L4). R1-R6s signal to LMCs synaptically, using inhibitory transmitter, histamine (Hardie, 1989), while the feedback synapses from LMCs to R1-R6s use excitatory transmitters (Davis et al., 2018). ***A***, R1-R6 rhabdomere has 30,000 microvilli. Each microvillus (inset) is a semi-independent photon sampling unit with full transduction reactions to stochastically absorb incoming photons and adaptively transduce them to quantum bumps (QBs). These reactions are modelled by 20 differential equations with 50 fixed parameters (no free parameters) using the Gillespie algorithm (Song et al., 2012). ***B***, Because each microvillus recovers from its previous QB within ~50-200 ms (refractoriness), its probability to convert its next absorbed photon (dot rows) to a QB increases in time from 0 to 1, with not every absorbed photon causing a QB (Song et al., 2012). ***C***, The photons follow Poisson statistics to sum up the light stimulus (green) and the QBs sum up the macroscopic light-induced current (LIC, blue). ***D***, LIC drives a HH R1-R6 membrane model (parameters in Tables 1–2), simulating a voltage response. ***E***, This simulation (gray) is compared to a real recording (purple; intracellular voltage signal to same light stimulus. See ***In vivo intracellular recordings***). ***F***, Flat (tonic) feedback conductance (gray; mimicking synaptic input from the LMCs) is injected into the R1-R6 model (with LIC) and its waveform is shaped dynamically (wine) in a closed-loop until the resulting simulated photoreceptor voltage response matches the real recording (***E***).

Modules 1-3 simulate the stochastic phototransduction cascade in the rhabdomere. Because the mutants’ phototransduction reactions were physiologically intact (as shown in the Results), all the parameters were fixed and kept the same in the simulations (50 parameters in 20 Equations); the mathematical details and parameters values are given in (Song et al., 2012; Juusola et al., 2015). Module 4 models the R1-R6 plasma membrane using deterministic continuous functions (HH model), in which parameters scale the model response to light stimulation, and now also to the estimated synaptic feedback (see below), approximating the recorded response (Figs 2*D-F*).

### Estimating excitatory synaptic feedback conductance

Differences between the simulated and recorded responses (Fig. 2*E*) should reflect a real photoreceptor’s synaptic feedback dynamics - input from LMCs (Zheng et al., 2006; Rivera-Alba et al., 2011; Dau et al., 2016), which the original R1-R6 model lacks (Juusola et al., 2017). Using these differences, one can work out the synaptic input current to R1-R6s.

The synaptic feedback current to each recorded R1-R6, whether wild-type or mutant, was extrapolated computationally by using the same fixed LIC (to the naturalistic light stimulus, Fig. 2*C*) with their specific *Shaker* (I_A_) and *Shab* (I_Ks_) current dynamics (Fig. 2*D*; see ***Whole-Cell Recordings***, above, and Tables 1–2). In this procedure, a new flat (Fig. 2*F*, gray) conductance, representing the missing synaptic input, was injected to the full wild-type, *dSK*^−^, *dSlo*^−^ or *dSK*^−^;;*dSlo*^−^ R1-R6 model. This conductance waveform was then shaped up in a closed-loop (Fig. 2*F*, wine), by our open-source Matlab software (GitHub access below), until the model’s voltage response matched the corresponding recorded voltage signal (a single R1-R6’s average response; see ***In vivo intracellular recordings***, above) for the same light stimulus.

**Table 1.**
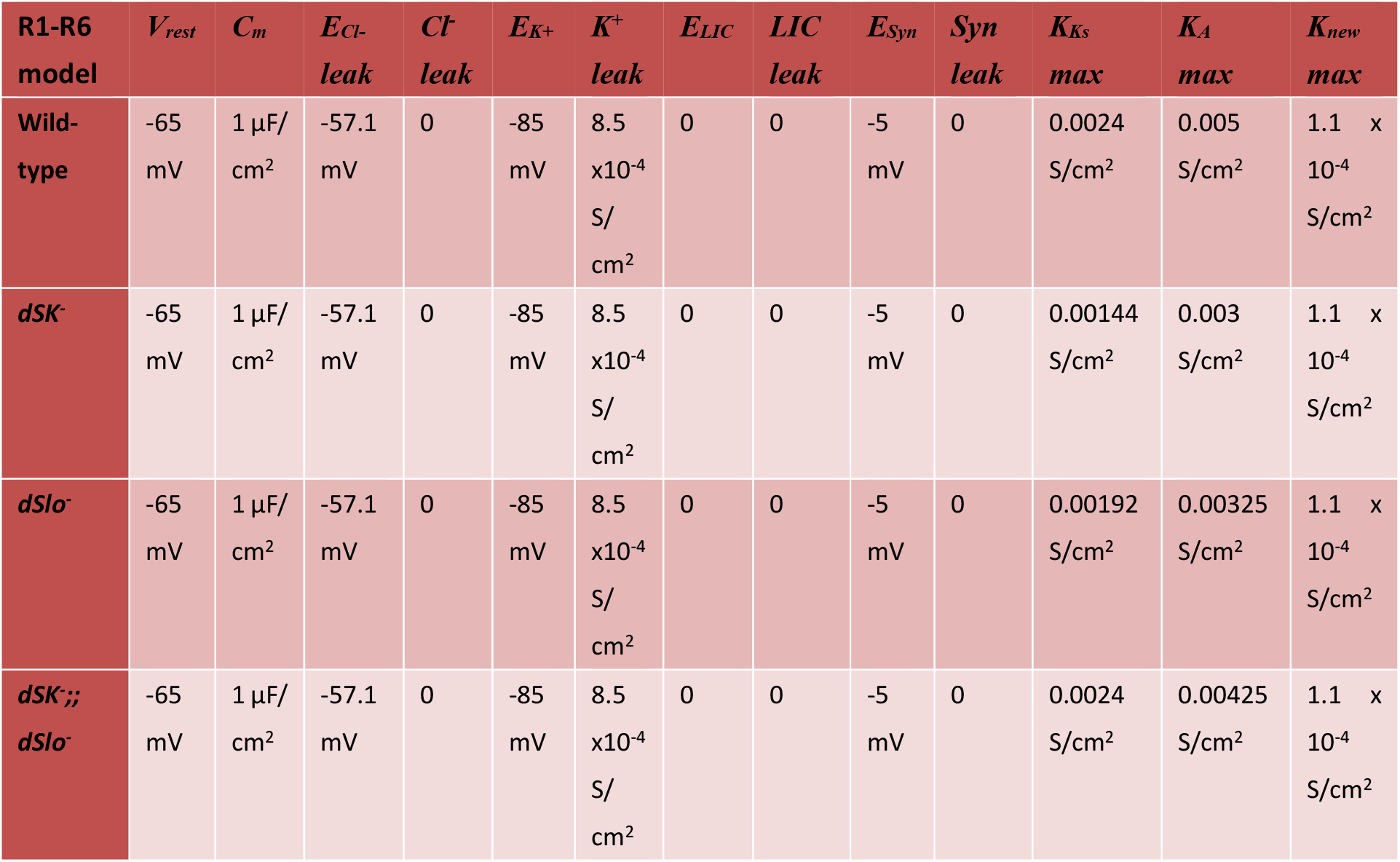
Conservative photoreceptor HH-membrane model parameters: Note, to be consistent: *Shab* K^+^-current is abbreviated as Ks and *Shaker* K^+^-current as A

**Table 2.**
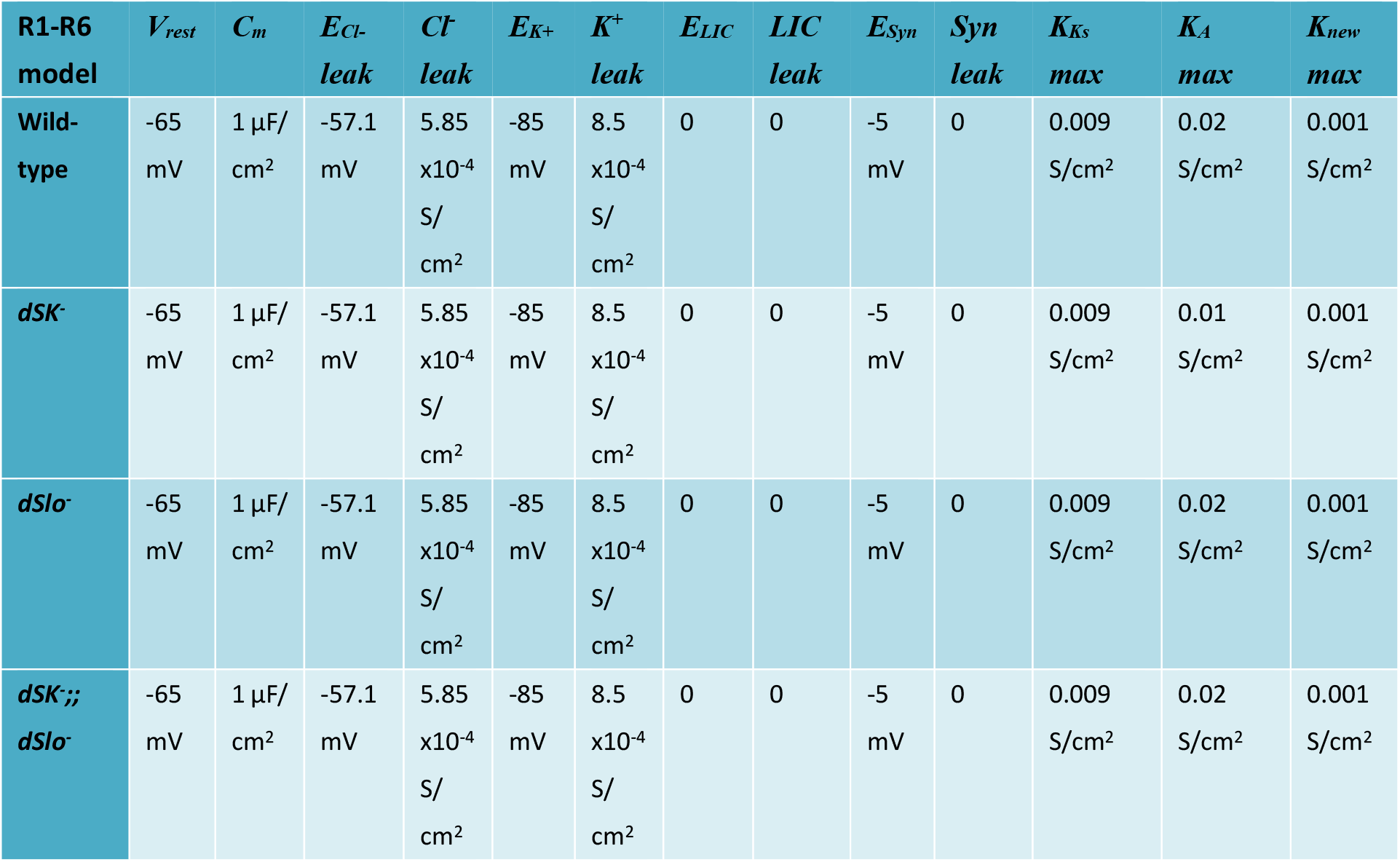
Speculative photoreceptor HH-membrane model parameters:

5 Table 2, Speculative photoreceptor HH-membrane model parameters:
| R1-R6<br>model | $V_{rest}$ | $C_m$ | $E_{Cl-}$<br>leak | $Cl^-$<br>leak | $E_{K+}$ | $K^+$<br>leak | $E_{LIC}$ | $LIC$<br>leak | $E_{Syn}$ | $Syn$<br>leak | $K_{Ks}$<br>max | $K_A$<br>max | $K_{new}$<br>max |
| --- | --- | --- | --- | --- | --- | --- | --- | --- | --- | --- | --- | --- | --- |
| Wild-type | -65<br>mV | 1 μF/<br>cm<sup>2</sup> | -57.1<br>mV | 5.85<br>x10<sup>-4</sup><br>S/<br>cm<sup>2</sup> | -85<br>mV | 8.5<br>x10<sup>-4</sup><br>S/<br>cm<sup>2</sup> | 0 | 0 | -5<br>mV | 0 | 0.009<br>S/cm<sup>2</sup> | 0.02<br>S/cm<sup>2</sup> | 0.001<br>S/cm<sup>2</sup> |
| *dSK*<sup>-</sup> | -65<br>mV | 1 μF/<br>cm<sup>2</sup> | -57.1<br>mV | 5.85<br>x10<sup>-4</sup><br>S/<br>cm<sup>2</sup> | -85<br>mV | 8.5<br>x10<sup>-4</sup><br>S/<br>cm<sup>2</sup> | 0 | 0 | -5<br>mV | 0 | 0.009<br>S/cm<sup>2</sup> | 0.01<br>S/cm<sup>2</sup> | 0.001<br>S/cm<sup>2</sup> |
| *dSlo*<sup>-</sup> | -65<br>mV | 1 μF/<br>cm<sup>2</sup> | -57.1<br>mV | 5.85<br>x10<sup>-4</sup><br>S/<br>cm<sup>2</sup> | -85<br>mV | 8.5<br>x10<sup>-4</sup><br>S/<br>cm<sup>2</sup> | 0 | 0 | -5<br>mV | 0 | 0.009<br>S/cm<sup>2</sup> | 0.02<br>S/cm<sup>2</sup> | 0.001<br>S/cm<sup>2</sup> |
| *dSK*<sup>-</sup>;;<br>*dSlo*<sup>-</sup> | -65<br>mV | 1 μF/<br>cm<sup>2</sup> | -57.1<br>mV | 5.85<br>x10<sup>-4</sup><br>S/<br>cm<sup>2</sup> | -85<br>mV | 8.5<br>x10<sup>-4</sup><br>S/<br>cm<sup>2</sup> | 0 | 0 | -5<br>mV | 0 | 0.009<br>S/cm<sup>2</sup> | 0.02<br>S/cm<sup>2</sup> | 0.001<br>S/cm<sup>2</sup> |

Remarkably, here, the predicted synaptic input from LMCs to a specific R1-R6 was extracted purely from the difference between the photoreceptor recording and simulation. Yet, it much resembled typical intracellular LMC voltage responses to the same light stimulus (see Results). This implies that the computationally extracted synaptic feedback, which systematically and consistently differed between the wild-type and mutant R1-R6s (see Results), would closely resemble the real excitatory feedback these cells receive from the lamina network *in vivo*. Such a strong logical agreement between these two independently obtained results highlights the explorative power of this new hybrid simulation/recording approach, validating its use.

Note, one cannot calculate the synaptic current from *ex vivo* dissociated cells, even if it was possible to retain their axon terminals, as these have different capacitance, input currents, extracellular milieu and voltage gradients than the *in vivo* photoreceptors. *In vivo*, the retina and lamina are partitioned by a glia-barrier, which keeps their respective extracellular fields at different potentials (Shaw, 1984).

### Estimating ATP Consumption for Information Transmission in Wild-type and Mutant R1-R6s

While the microvilli, which form the photosensitive R1-R6 rhabdomere (Fig. 2*B*), generate the LIC, the photo-insensitive plasma membrane uses many voltage-gated ion channels to adjust the LIC-driven voltage responses. In response to LIC, these open and close, regulating the ionic flow across the plasma membrane and further modulating tonic neurotransmitter (histamine) release at the photoreceptor-LMC synapse (Hardie, 1989; Uusitalo et al., 1995b). In return, tonic excitatory synaptic feedback from the LMCs participate in shaping the R1-R6 voltage response (Fig. 2*F*) (Zheng et al., 2006; Zheng et al., 2009; Dau et al., 2016). But to maintain the pertinent ionic concentrations in- and outside, R1-R6s rely upon other proteins, such as ion cotransporters, exchangers and pumps, to uptake or expel ions. The work of moving ions against their electrochemical gradients consumes energy (ATP), and a R1-R6’s ATP consumption thus much depends on the ionic flow dynamics through its ion channels (Laughlin et al., 1998). To approximate these dynamics during light responses, we used our HH R1-R6 body model (Niven et al., 2003; Song et al., 2012), which models the ion channels as conductances.

The HH model has these ion transporters: 3Na^+^/2K^+^-pump, 3Na^+^/Ca^2+^-exchanger and Na^+^/K^+^/2Cl^−^ mechanisms to balance the intracellular ionic fluxes. Na^+^/K^+^/2Cl^−^ cotransporter balances with the voltage-dependent Cl^−^ and Cl^−^ leak conductances, maintaining intracellular Cl^−^-concentration. Ca^2+^ influx in the LIC (~41%) is then expelled by 3Na^+^/Ca^2+^-exchanger in 1:3 ratio in exchange for Na^+^ ions. Although there is K^+^ influx in LIC (~24%), this is not enough to compensate K+ leakage through voltage-gated K^+^ conductances and K^+^ leaks. Apart from a small amount of K^+^ intake through Na^+^/K^+^/2Cl^−^-cotransporter, 3Na^+^/2K^+^-pump is the major K^+^ uptake mechanism. It consumes 1 ATP molecule to uptake 2 K^+^ ions and extrudes 3 Na^+^ ions. Because it is widely regarded as the major energy consumer in the cell, we use only the pump current (*I_p_*) to estimate the ATP consumption (Skou, 1965; Laughlin et al., 1998; Skou, 1998). For these estimates, we generated two separate photoreceptor membrane models: a conservative one (Table 1; containing the known voltage-sensitive and leak potassium conductances) and a speculative one (Table 2; by adding an unconfirmed chloride conductance and leak, now balanced with larger voltage-sensitive K^+^ conductances). Their differences helped us to work out how the earlier proposed hypothetical homeostatic compensation through leak- or chloride channel expression (Niven et al., 2003; Vähäsöyrinki et al., 2006) would change a photoreceptor’s ATP consumption.

From the equilibrium of K^+^ fluxes, *I_p_* can be calculated as follows:

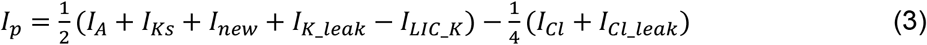

where *I_A_*, *I_Ks_*, *I_new_*, and *I_K_leak_* are the currents through *Shaker, Shab*, new, and K_leak channels, respectively, *I_LIC_K_* is the K^+^ influx in LIC and *I_Cl_* and *I_Cl_leak_* are the currents through the voltage-gated Cl^−^ and Cl^−^ leak channels, respectively. These currents can be calculated from the reverse potential of individual ions and their HH model produced conductances using Ohm’s law:

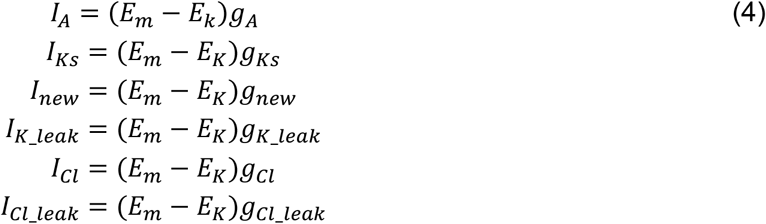

Using *I_p_*, the number of ATP molecules hydrolyzed per second can be calculated:

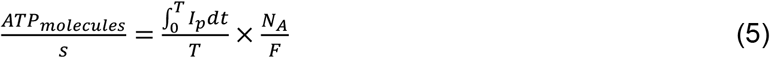

where *N_A_* is Avogadro’s constant and *F* is Faraday’s constant. The ATP usage per bit of information was calculated by dividing the estimated ATP molecules hydrolyzed in 1 s by the estimated information transfer rates (bits/s). We did not model the respective pump dynamics because, for the purpose of calculating ATP, only the time-integrated ionic fluxes count, not the time constants.

Previously, because of lack of a complete model for the photosensitive membrane, the LIC has only been estimated at the steady-state, or DC (Laughlin et al., 1998; Niven et al., 2007), when the sum of all currents across the model membrane equals zero:

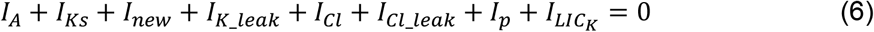

Thus here, the conservative photoreceptor membrane model (Table 1) lacked *I_Cl_leak_* and *I_Cl_* in Eqs. 3, 4 and 6, whereas the speculative model (Table 2) included them. But for both membrane models, because we estimated LIC directly from the stochastic phototransduction model (above), we could calculate a R1-R6’s energy cost in response to any arbitrary light pattern, including naturalistic stimulation. Thus, our phototransduction cascade model provides the *functional* equivalence to the light-dependent conductance used in the previously published steady-state models (Laughlin et al., 1998; Niven et al., 2007).

Finally, we note that if the ion transporters, including 3Na^+^/2K^+^-pump, were inherently noisy, their metabolic work would reduce a photoreceptor’s information transfer rate. However, such effects are likely very small as the recorded and simulated photoreceptor information transfer rate estimates effectively match over a broad range of light stimuli (Juusola et al., 2017). Conversely, in concordance the data processing theorem (Shannon, 1948; Juusola and de Polavieja, 2003; Cover and Thomas, 2006), ion transporters cannot increase a photoreceptor’s information transfer rate as they cannot increase sample (QB) rate changes that sum up the response, increasing its *SNR*(*f*).

### Code and Software Accessibility

Biophysical *Drosophila* model (Matlab) is freely available from github (https://github.com/JuusolaLab/Microsaccadic_Sampling_Paper/tree/master/BiophysicalPhotoreceptorModel). LMC feedback to R1-R6 estimation (Matlab) is freely available from github (https://github.com/JuusolaLab/SK_Slo_Paper/tree/master/LMCFeedbackToR1-R6Model).

Information estimation (Matlab) is freely available from github (https://github.com/JuusolaLab/Microsaccadic_Sampling_Paper/tree/master/SNRAnalysis).

### Histology

#### Electron Microscopy

3-to-7-day-old dark/light-reared *Drosophila* were cold anesthetized on ice and transferred to a drop of pre-fixative [modified Karnovsky’s fixative: 2.5% glutaraldehyde, 2.5% paraformaldehyde in 0.1 M sodium cacodylate buffered to pH 7.3 – as per (Shaw et al., 1989)] on a transparent agar dissection dish. Dissection was performed using a shard of a razor blade (Feather S). Flies were restrained on their backs with insect pins through their lower abdomen and distal proboscis. Their heads were severed, proboscis excised, and halved. The left half-heads were collected in fresh pre-fixative and kept for 2 h at room temperature (21 ± 1 °C) under room light.

After pre-fixation, the half-heads were washed (2 × 15 min) in 0.1 M Cacodylate buffer, and then transferred to a 1 h post-fixative step, comprising Veronal Acetate buffer and 2% Osmium Tetroxide in the fridge (4°C). They were moved back to room temperature for a 9 min wash (1:1 Veronal Acetate and double-distilled H_2_O mixture), and serially dehydrated in multi-well plates with subsequent 9 min washes in 50, 70, 80, 90, 95, and 2 × 100% ethanol. Post-dehydration, the half-heads were transferred to small glass vials for infiltration. They were covered in Propylene Oxide (PPO) for 2 × 9 min, transferred into a 1:1 PPO:Epoxy resin mixture (Poly/Bed^®^ 812) and left overnight. The following morning, the half-heads were placed in freshly made pure resin for 4 h, and placed in fresh resin for a further 72 h at 60 °C in the oven. Fixation protocol was provided by Professor Ian Meinertzhagen (Dalhousie University, Canada).

Embedded half-heads were first sectioned (at 0.5 μm thickness) using a glass knife, mounted in an ultramicrotome (Reichert-Jung Ultracut E, Germany). Samples were collected on glass slides, stained using Toluidine Blue and observed under a light microscope. This process was repeated and the cutting angle was continuously optimized until the correct orientation and sample depth was achieved; stopping when approximately 40 ommatidia were discernible. The block was then trimmed and shaped for ultra-thin sectioning. The trimming is necessary to reduce cutting pressure on the sample-block and resulting sections, thus helping to prevent “chattering” and compression artifacts.

Ultra-thin sections (85 nm thickness) were cut using a diamond cutting knife (DiATOME Ultra 45°, USA), mounted and controlled using the ultramicrotome. The knife edge was first cleaned using a polystyrol rod to ensure integrity of the sample-blocks. The cutting angles were aligned and the automatic approach- and return-speeds set on the microtome. Sectioning was automatic and samples were collected in the knife water boat. Sections were transferred to Formvar-coated mesh-grids and stained for imaging: 25 min in Uranyl Acetate; a double-distilled H_2_O wash; 5 min in Reynolds’ Lead Citrate (Reynolds, 1963); and a final double-distilled H_2_O wash.

#### Conventional microscopy

Heads of 8-day-old dark/light-reared female and male flies were bisected, fixed, and embedded as explained previously (Chinchore et al., 2009). 1 μm eye cross sections were cut using a Sorvall ultra microtome MT-1 (Sorvall, CT), stained with toluidine blue, and inspected using a Zeiss Axioplan2 microscope. Digital images were taken using Optronics DEI-750 camera (Optronics) and MetaVue (Universal Imaging) software.

#### Experimental Design and Statistical Analysis

Figures show mean ± SD (or SEM) for each group (wild-type, *dSK^−^, dSlo*^−^ and *dSK*^−^;;*dSlo*^−^), and typically also the individual values for each recordings (marked as ○). Significance between two groups was calculated using 2-tailed paired Student’s t-test (both with and without the equal variance assumption). In Figs 1 and 4, we used also one-way ANOVA for multiple comparisons between each group. For the analyses, we used OriginLab 2018b, Graphpad Prism v5 and Matlab statistical toolbox. Specific p-values and sample sizes are indicated in the relevant figures and/or legends.

## Results

### Absence of dSK and dSlo Shapes Photoreceptor Responses

To examine how Ca^2+^-activated K^+^ channels shape *Drosophila* photoreceptor voltage output, we performed *in vivo* intracellular recordings (Fig. 3*A*) from R1-R6 somata (Fig. 3*B*) in the retinae of *dSlo^−^, dSK*^−^ and *dSK*^−^;;*dSlo*^−^ null mutants and wild-type flies, using conventional sharp microelectrodes. Briefly dark-adapted (~20 s) mutant R1-R6s responded to logarithmically brightening light flashes with increasing graded depolarizations (Fig. 3*C*), having wild-type-like or slightly smaller amplitudes (Fig. 3*D*). However, both and *dSK*^−^ and *dSK*^−^;;*dSlo*^−^ R1-R6 outputs peaked faster (Fig. 3*E*; mean time-to-peak) and decayed earlier (Fig. 3*F*; mean half-width) to their respective resting potentials than the wild-type. While those of *dSlo*^−^ R1-R6s, in contrast, showed decelerated dynamics, lasting longer than the wild-type except at the highest intensities (Figs 3*C* and 3*F*).

**Figure 3.**
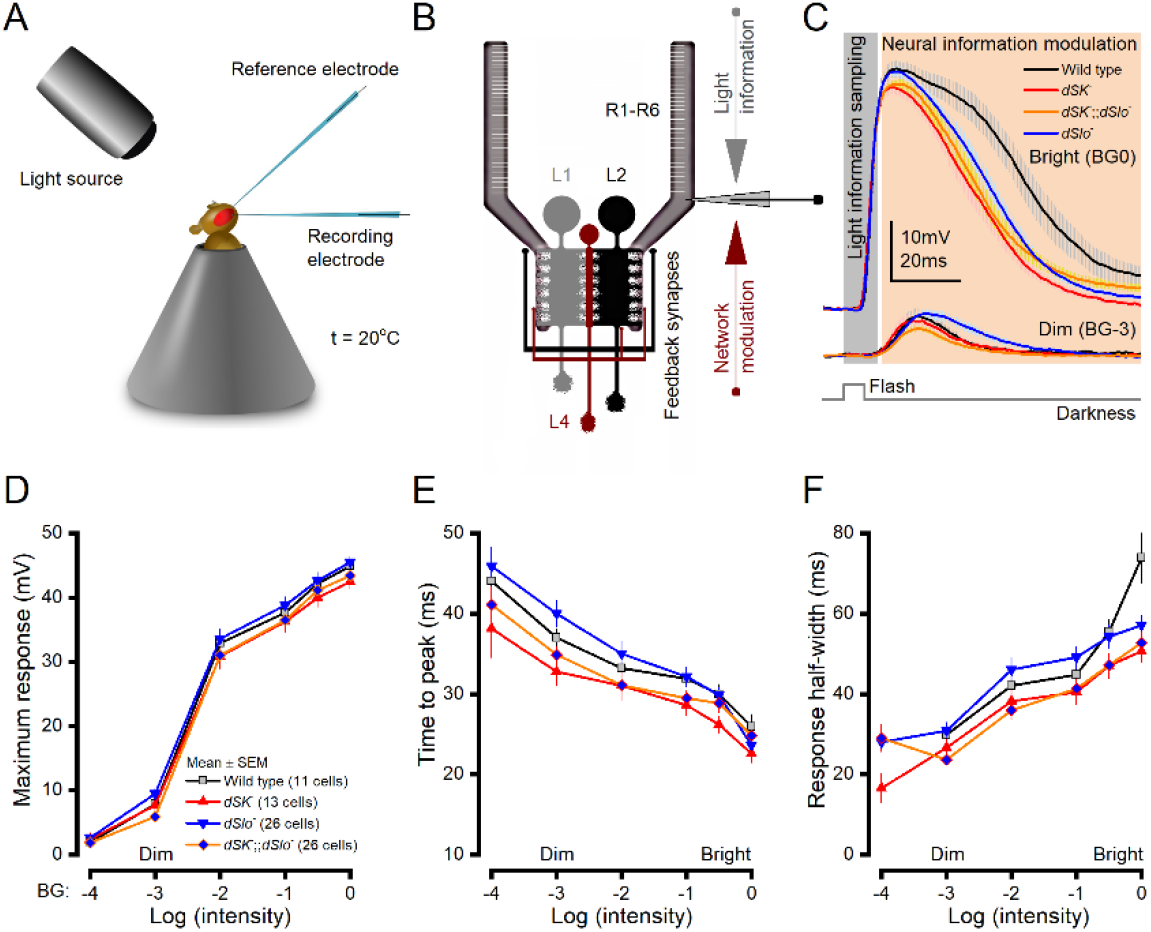
R1-R6 Photoreceptors of Different Ca^2+^-Activated K^+^ Channel Null-Mutants Show Distinctive Response Dynamics to Light Flashes. ***A***, Recordings were performed *in vivo* from R1-R6 somata using conventional sharp microelectrodes. ***B***, 30,000 *microvilli*, which form a R1-R6’s light-sensor, the rhabdomere (comb-like structure), sample light information (incoming photon rate changes). R1-R6 axon terminals then transmit these signals to the lamina network through sign-inverting (histaminergic) output synapses (to L1-L3 monopolar cells and amacrine cells) and receive synaptic feedback (network modulation) in return (Meinertzhagen and O’Neil, 1991; Rivera-Alba et al., 2011); the schematic highlights excitatory feedbacks from L2, lamina intrinsic amacrine neurons (Lai) and L4 to photoreceptor terminals. Because R1-R6s are short and have large length constants, synaptic feedback also influences their somatic response waveforms (Zheng et al., 2006; Dau et al., 2016). ***C***, The average *dSlo*^−^, *dSK*^−^, *dSK*^−^;;*dSlo*^−^ and wild-type responses to 10 ms bright and dim flashes. Their corresponding delay and rise times (≤20 ms from the flash onset; light gray area) were similar, suggesting intact light information sampling. But the mutant R1-R6 responses decayed (in light brown area) either faster or slower than the wild-type, suggesting differences in neural tuning. ***D***, Mutant and wild-type R1-R6 responses had comparable maximum amplitudes over the tested flash intensity range, resulting in similar V/log(I) saturation curves. ***E***, *dSK*^−^;;*dSlo*^−^ and *dSK*^−^ responses peaked, on average, sooner than the wild-type to all test intensities, with significantly shorter *dSK*^−^ values for BG-0.5 (p = 0.028) and BG-1 (p = 0.047). Conversely, *dSlo*^−^ responses peaked later than the wild-type to all but the two brightest flashes. Moreover, these responses peaked significantly later than those of *dSK*^−^ at BG-0.5 (p = 0.035) and BG-3 (p = 0.013), and *dSK*^−^;;*dSlo*^−^ at BG-2 (p = 0.038) and BG-3 (p = 0.013). ***F***, Wild-type response half-widths to the brightest flash (BG0) were significantly longer than those of *dSK*^−^ (p = 0.003) and *dSK*^−^;;*dSlo*^−^ (p = 8.56 × 10^−4^) R1-R6s. Conversely, *dSlo*^−^ responses, on average, lasted the longest over a broad flash intensity range; vs. *dSK*^−^;;*dSlo*^−^: at BG-1 (p = 0.042) BG-2 (p = 0.011) and BG-3 (p = 0.007). ***D-F***: Mean ± SEM, two-tailed t-test.

Notably, however, in all the corresponding recordings, the early light-induced depolarizations (Fig. 3*C*; light grey area) were similar, implying that the mutant R1-R6s sampled light information normally. Thus, phototransduction reactions inside a R1-R6’s ~30,000 microvilli (photon sampling units; Fig. 3*B*), which form its light-sensor, the rhabdomere (Hardie and Juusola, 2015), seemed unaffected by the absence of Ca^2+^-activated K^+^ channels. But, instead, these mutant genotypes influenced more the subsequent neural information modulation phase (Fig. 3*C*; light brown area).

### Response Differences not from Homeostatic Ion Channel Expression

If a R1-R6 photoreceptor was an isolated system, missing Ca^2+^-activated K^+^-conductances would directly increase its membrane resistance, *R_m_*, and consequently its time constant (*τ_m_* = *R_m_*·*C_m_*; *C_m_* is membrane capacitance). This would slow down voltage responses to light changes. However, *in vivo*, as each R1-R6 features complex bioelectric interactions within its membrane and with its neural neighbors, the mutant responses showed far more sophisticated dynamics (Fig. 3), presumably reflecting homeostatic changes in these interactions (Marder and Goaillard, 2006; Vähäsöyrinki et al., 2006). Therefore, to work out what made the mutant R1-R6 outputs differ, we analyzed changes both in their intrinsic (membrane) properties and extrinsic (synaptic) feedback from the surrounding network.

We first asked whether the differences in *dSlo^−^, dSK*^−^ and *dSK*^−^;;*dSlo*^−^ R1-R6 voltage responses resulted from homeostatic somatic conductance changes. These would affect their membrane resistances, accelerating or decelerating signal conduction. For example, missing dSK channels in *dSK*^−^ photoreceptors could be compensated by up-regulating dSlo channel expression, for which these cells carry a normal gene; and *vice versa* in *dSlo*^−^ photoreceptors. Alternatively, the cells could increase K^+^- or Cl^−^-leak-conductances (Niven et al., 2003; Vähäsöyrinki et al., 2006). While such intrinsic homeostatic mechanisms could accelerate *dSK*^−^ R1-R6 output, these would also lower their resting potentials; by reducing depolarizing Ca^2+^-load and/or increasing hyperpolarizing K^+^/Cl^−^ loads. Equally, a lack of such homeostatic ion channel expression changes could have contributed to *dSlo*^−^ photoreceptors’ slower signaling.

To test these hypotheses, we measured *in vivo* somatic electrical membrane properties in dark-adapted mutant and wild-type R1-R6s (Fig. 4*A*) using single-electrode current-clamp (e.g. Juusola and Weckström, 1993). We found that all the mutant R1-R6s charged smaller, but broadly wild-type-like voltage responses to injected current pulses (Fig. 4*B*). Depolarization to positive currents showed characteristic outward rectification (arrows), caused by activation of voltage-dependent K^+^ channels (Hardie, 1991a; Hardie et al., 1991; Juusola and Hardie, 2001b; Vähäsöyrinki et al., 2006), while hyperpolarization to negative currents, in effect, charged their membranes passively.

**Figure 4.**
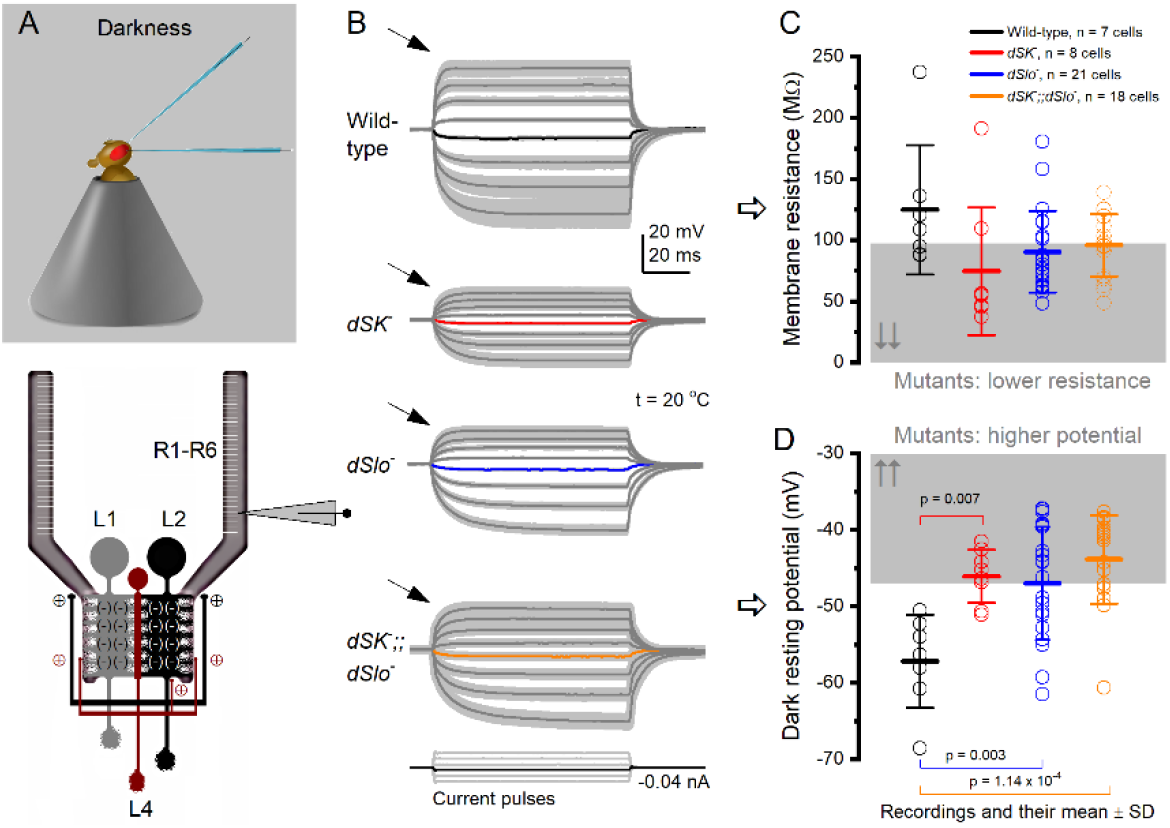
In Darkness, R1-R6 Photoreceptors of Ca^2+^-Activated K^+^ Channel Null-Mutants Have Lower Membrane Resistances and Higher Resting Potentials. ***A***, *In vivo* R1-R6 recordings. R1-R6 terminals provide histaminergic feedforward inhibition to LMCs. In return, R1-R6s receive excitatory feedback from L2 and L4 monopolar cells. ***B***, Voltage responses of dark-adapted wild-type and mutant R1-R6s to intracellular current pulse injections. The arrows indicate outward rectification; caused by voltage-sensitive *Shaker* and *Shab* K^+^-conductance activation to fast membrane depolarizations. ***C***, Mean wild-type R1-R6 input resistance (124.8 ± 52.7 MΩ, n = 7 cells) is significantly higher than that for all the mutant recordings (89.8 ± 34.5 MΩ, n = 47, p = 0.024), but not for each mutant-type separately (*dSK*^−^, 74.6 ± 52.0 MΩ, p = 0.071, n = 8; *dSlo*-, 90.4 ± 33.4 MΩ, p = 0.232, n = 21; *dSK*^−^;;*dSlo*^−^, 95.9 ± 25.6 MΩ, p = 0.519, n = 18). ***D***, Mutant R1-R6s are more depolarized than the wild-type photoreceptors in darkness (wild-type, −57.2 ± 6.1 mV; *dSK*^−^, −46.1 ± 3.5 mV; *dSlo*-, −47.0 ± 7.4 mV; *dSK*^−^;;*dSlo*^−^, - 43.9 ± 5.8 mV). ***C-D***: Mean ± SD, p-values from ANOVA for multiple comparisons with Bonferroni-test.

The membrane input resistances of the mutant R1-R6s (Fig. 4*C*), as determined by small hyperpolarizing responses to −0.02 nA current steps, were characteristically lower than in the wild-type (Juusola and Hardie, 2001b; Niven et al., 2003), with the mean resistance of *dSK*^−^ R1-R6s being the lowest (*cf*. Abou Tayoun et al., 2011). Most crucially, however, the mutant (dSK^−^, *dSlo*^−^ and *dSK*^−^;;*dSlo*^−^) photoreceptors’ resting potentials (Fig. 4*D*), instead of being more hyperpolarized, were >10 mV more depolarized than the wild-type. Here, if *dSK*^−^ or *dSlo*^−^ R1-R6s’ intrinsic signaling properties were regulated homeostatically, by ion channel expression (as hypothesized), then their resting potential in darkness should have been below the wild-type range, rather than above it. Also, the higher resting potentials (Fig. 4*D*) and lower membrane resistances (Fig. 4*C*) should have accelerated signal conduction. Yet, the mean *dSlo*^−^ R1-R6 voltage response time-to-peak values to intermediate light flash intensities were, in fact, slower than in the wild-type (Figs 3*E* and 3*F*).

Hence, collectively, these results suggested that the accelerated (*dSK*^−^ and *dSK*^−^;;*dSlo*^−^) and decelerated (*dSlo*^−^) light-induced voltage response dynamics of the mutant photoreceptors (Figs 3*B* and 3*C*) unlikely resulted from compensatory expression of leak- or Ca^2+^-activated K^+^ channels at the somata, but required other/further mechanisms. Notice, however, these results were recorded from dark-adapted R1-R6s at relative rest; without light-induced conductance (LIC) interactions. During brightening light stimulation, LIC and other conductances increase progressively with membrane depolarization (e.g. Song et al., 2012; Juusola et al., 2017), reducing resistance further by >> 10-fold (e.g. Juusola and Weckström, 1993), as our recordings and simulations clarify later on.

### Response Differences not by Transduction or K^+^ Conductance Differences

To eliminate the possibility that developmental morphological defects in the mutant R1-R6s would have caused their altered responses, we assessed the mutant and wild-type eyes/retinae using both electron- (Fig. 5*A*, above) and light-microscopy (below). We found no obvious morphological differences between the eyes; with each method displaying highly ordered ommatidia with normal looking intact R1-R7 photoreceptor rhabdomeres.

**Figure 5.**
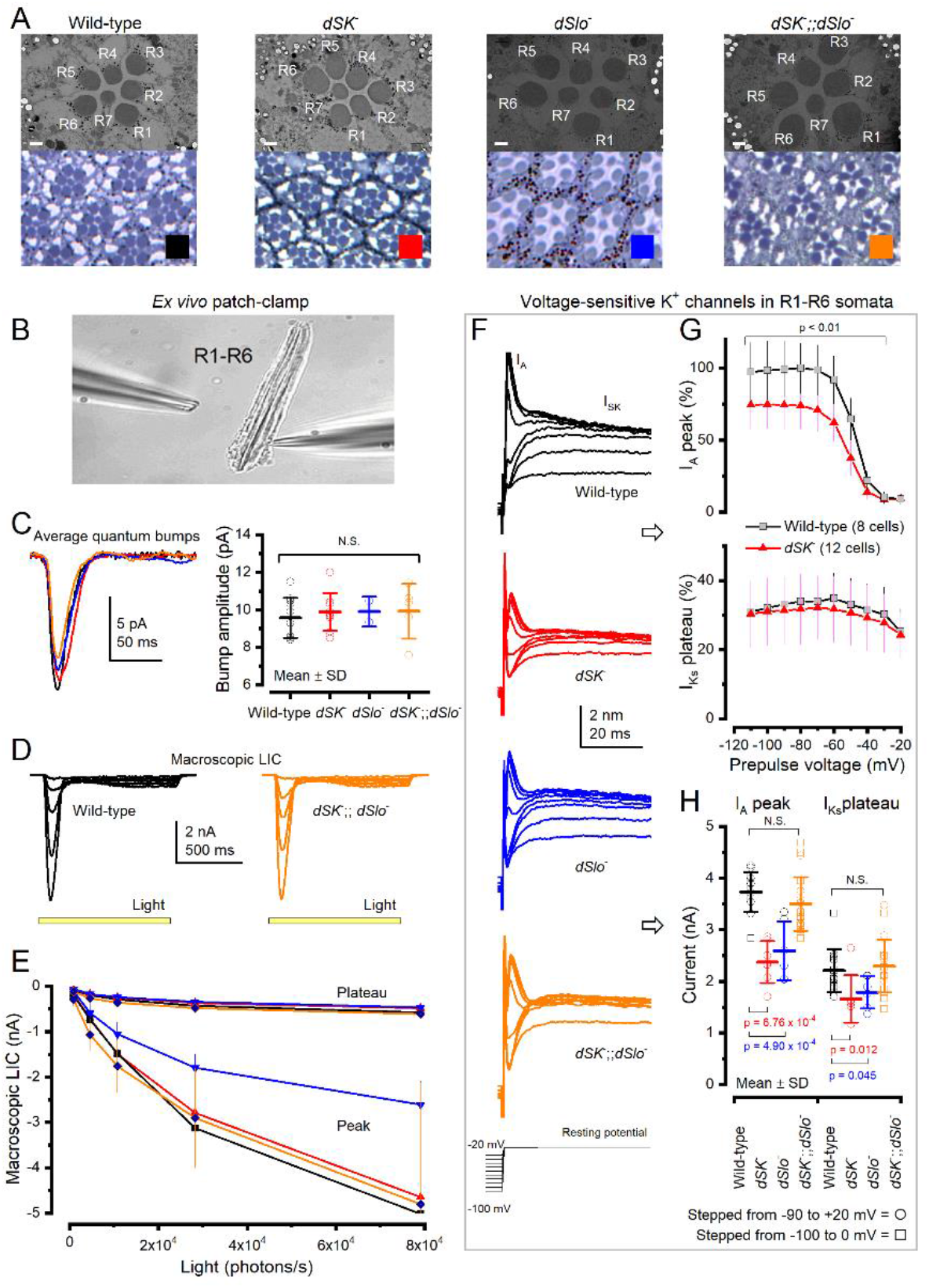
*dSK*^−^, *dSlo*^−^ and *dSK*^−^;;*dSlo*^−^ Photoreceptors Have Normal Morphology, Light-Induced Currents (LIC) but 8-40% Reduced Light-Insensitive I_A_ Currents. ***A***, The mutant retinae appear structurally intact, with R1-R7s having wild-type-like rhabdomeres and pigmentation; 1 μm EM scale bars. ***B***, Whole-cell recordings were performed from dissociated ommatidia. ***C***, Mutant and wild-type R1-R6 quantum bump (QB) waveforms and their amplitude distributions to dim flashes are similar. ***D***, Wild-type and *dSK*^−^;;*dSlo*^−^ R1-R6 LIC responses to 1 s light pulses of different brightness. ***E***, Macroscopic LIC peak and plateau responses were similar, with the normal experimental variation, indicating that dSK and dSlo deletions do not affect phototransduction. The smaller *dSlo*^−^ max LIC probably resulted from these ommatidia being smaller, reflecting *dSlo*^−^ mutants reduced yield/health. ***F***, Wild-type and mutant R1-R6s’ voltage-sensitive outward K^+^ currents to increased voltage steps contain both the transient *Shaker* (I_A_) and sustained delayed rectifier, *Shab*, (I_KS_) components. ***G***, *dSK*^−^ R1-R6 K^+^-currents have a reduced I_A_ but near normal-sized I_KS_. ***H***, On average, the maximum I_A_ and I_KS_ currents in *dSK*^−^ and *dSlo*^−^ R1-R6s are a bit smaller than the wild-type (*dSK*^−^ I_A_: 36.4% < wild-type, I_Ks_: 24.9% < wild-type; *dSlo*^−^ I_A_: 30.6% < wild-type, I_Ks_: 19.0% < wild-type) but wild-type-like in *dSK*^−^;;*dSlo*^−^ R1-R6s.

Nevertheless, deletion of dSlo, dSK or both could still affect intracellular [Ca^2+^] regulation, and thus potentially alter microvillar phototransduction functions indirectly (Song et al., 2012; Hardie and Juusola, 2015), modifying sampling, amplification or integration of light-induced currents (LIC). We, therefore, used whole-cell recordings in dissociated ommatidia (Hardie, 1991b) (Fig. 5*B*) to compare the mutant and wild-type R1-R6s’ elementary responses (quantum bumps, QBs) to single photons (Fig. 5*C*) and macroscopic LICs to light pulses (Figs 5*D* and 5*E*). In this preparation, photoreceptor axon terminals were severed, cutting off any synaptic feedback from the lamina network to R1-R6s (Zheng et al., 2006).

We found the mutant R1-R6s’ bump amplitudes and waveforms (Fig. 5*C*) and macroscopic LICs (Figs 5*D* and 5*E*) to increasing light intensities wild-type-like, showing normal dynamics within the normal experimental variation. Here, the smaller *dSlo*^−^ LIC maxima likely resulted from the smaller size of these homozygotic mutant flies due to their lower yield/reduced health. Thus, deletion of dSlo, dSK or both channels neither disrupted the microvillar R1-R6 morphology nor its phototransduction functions, again suggesting that the mutant R1-R6s would sample light information like their wild-type counterparts (see: Song et al., 2012; Hardie and Juusola, 2015; Juusola and Song, 2017).

Intriguingly, however, K^+^ conductances in dissociated *dSK*^−^ and *dSlo*^−^ R1-R6s showed slightly reduced (19-36%) fast A- (*I_A_* or *Shaker*) and delayed rectifier currents (*I_KS_* or *Shab*) (Figs 5*F-H*), while these currents were broadly wild-type-like in *dSK*^−^;;*dSlo*^−^ R1-R6s. The decrease in the *I_A_* and *I_Ks_* currents together with dSK or dSlo current removal should, with other things being equal, increase membrane resistance and its time constant, leading to slower voltage responses. Instead *in vivo*, we found resistance in all the mutant R1-R6s below the wild-type (Fig. 4*C*), with both *dSK*^−^ and *dSK*^−^;;*dSlo*^−^ R1-R6s responding faster and only *dSlo*^−^ R1-R6s slower (Fig. 3*E*), implying that homeostatic changes in K^+^ channel expression alone cannot explain their response differences.

Together, the observed normal rhabdomere morphology, wild-type-like LIC dynamics and only partly reduced photo-insensitive membrane conductances implied that the mutant R1-R6s’ accelerated or decelerated voltage responses, higher resting potentials and lower membrane resistance *in vivo* could not be induced by homeostatic ion channel expression changes in photoreceptor somata alone. But this would more require network adaptation (Nikolaev et al., 2009; Zheng et al., 2009), parallel changes in the synaptic network activity. In such scenarios, missing one or both Ca^2+^-activated K^+^ channels would cause a homeostatic (automatic) rebalancing of the bidirectional signal transfer between photoreceptor axon terminals and the lamina interneurons (Shaw, 1984; Zheng et al., 2006; Zheng et al., 2009; Abou Tayoun et al., 2011; Dau et al., 2016).

### dSK or dSlo Absence Changes Network Adaptation

In the adult *Drosophila* brain, dSlo and dSK share similar expression patterns with higher expression in the lamina and medulla neuropils and weaker in the retina (Becker et al., 1995; Abou Tayoun et al., 2011). Thus, theoretically, dSlo and dSK could co-participate in shaping the bidirectional signal transfer between R1-R6 photoreceptor axons and LMCs, which form columnar R-LMC-R network processing units in the lamina (Nikolaev et al., 2009; Zheng et al., 2009). Here, the deletion of one or the other ion channel could disrupt this balance.

We, therefore, next asked how Ca^2+^-activated K^+^ channels might contribute to network adaptation in the R-LMC-R system. We recorded *dSK^−^, dSlo^−^, dSK*^−^;;*dSlo*^−^ and wild-type R1-R6 responses to a repeated 1 s naturalistic light intensity time series stimulus (NS) (van Hateren, 1997) *in vivo*, and found each of them adapting differently (Fig. 6*A*).

**Figure 6.**
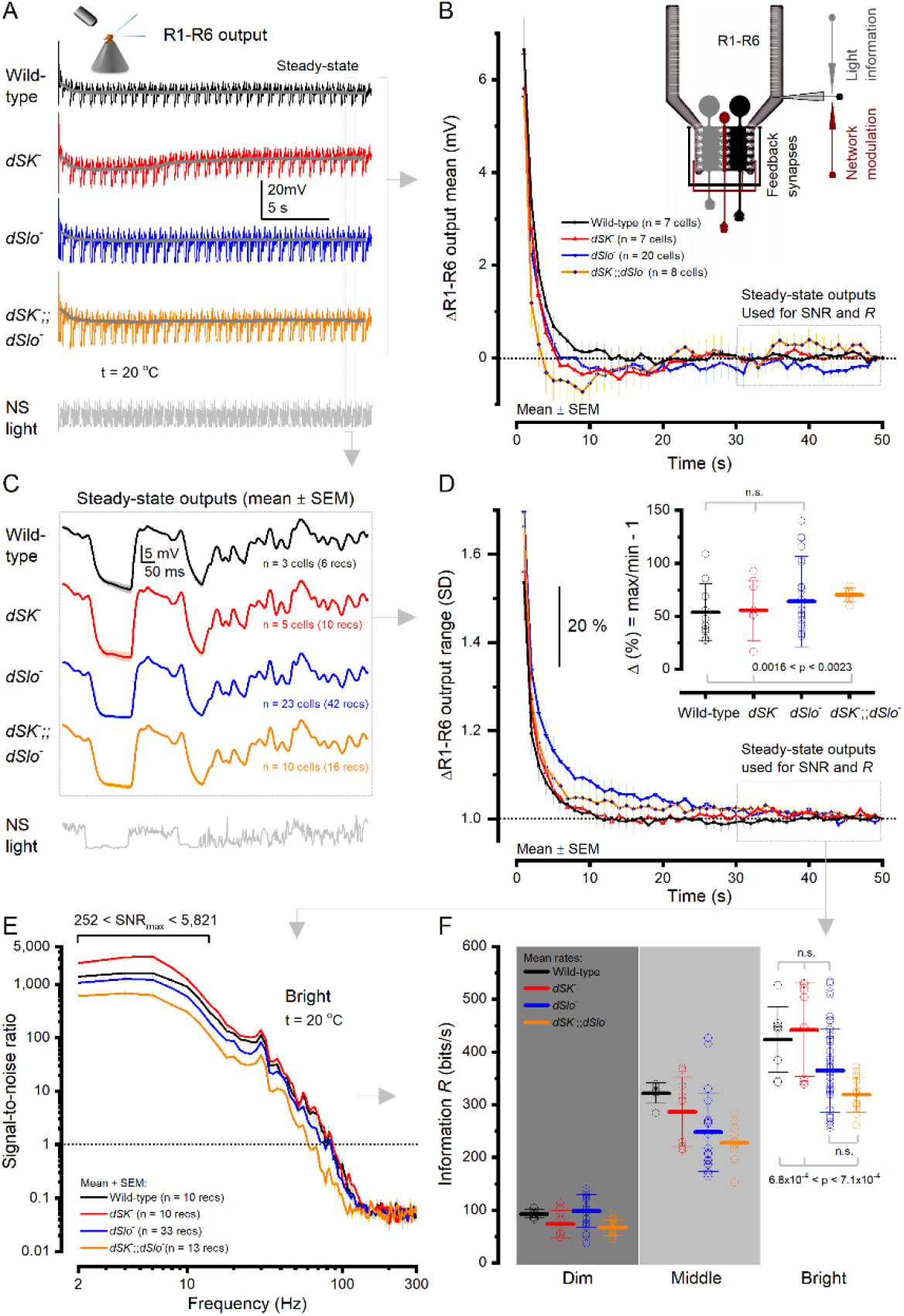
Adaptation Dynamics and Information Rates of Wild-Type and Ca^2+^-Activated K^+^ Channel Null-Mutant R1-R6 Photoreceptors. ***A***, Intracellular voltage responses to a repeated 1-s-long bright naturalistic light intensity time series stimulus (NS). ***B***, Change in the response mean (± SD) over the repeated stimulation. Mean wild-type and *dSlo*^−^ R1-R6 outputs declined near exponentially to steady-state, whereas adaptation in mean *dSK*^−^ and *dSK*^−^;;*dSlo*^−^ R1-R6 outputs depicted unique undershoots. ***C***, Mean waveforms ± SD of steady-state adapted 1 s responses. ***D***, Relative change in R1-R6 output range, measured as the standard deviation (SD) of the responses at each s during 50 s of stimulation (mean ± SD). Wild-type and *dSlo*^−^ R1-R6s desensitized during repeated stimulation, following exponential time constants. Wild-type R1-R6 output range contracted from 114% to 100% in about 26 s (*T*_wild-type_ = 1.96 ± 0.39 s); *dSlo*^−^ from 134% in about 19 s (*T*_dSlo4_ = 8.3 ± 1 s); *dSK*^−^;;*dSlo*^−^ from 133% in 8 s (*T*_dSK;;dSlo4_ = 3.4 ± 0.4 s). *dSK*^−^ and *dSK*^−^;;*dSlo*^−^ R1-R6s’ output ranges showed further sensitizing trends, reaching a steady-state after ~40 s. ***E***, Mutant and wild-type R1-R6s’ average signal-to-noise ratios, measured from their steady-state outputs to bright NS, are high and broadly similar. ***F***, Mutant and wild-type R1-R6s sampled information from dim, moderately intense (middle) and bright naturalistic stimulation in a comparable manner (mean ± SD; n = 10-33 recordings). *dSK*^−^;;*dSlo*^−^ R1-R6s had a marginally lower mean information transfer rate than the other genotype photoreceptors. In each genotype, R1-R6 information rates to the given stimulation vary naturally (up to ~200 bits/s) as each cell receives different amount of information from the network (Juusola et al., 2017).

The mean of the wild-type response (Fig. 6*B*, black trace; measured at each second) decreased approximately exponentially as the cells adapted to NS (Fig. 6*C*), reaching a relative steady-state in 15-20 s (Figs 6*B* and 6*C*). In contrast, the corresponding means of the mutant responses declined faster but then displayed unique genotype-specific undershooting. The means of *dSK*^−^ (red trace) and *dSK*^−^;;*dSlo*^−^ (orange) responses first decreased to their minima in <10 s, and then increased, as the cells gradually further depolarized, reaching a relative steady-state in 35-40 s; ~20 s later than the wild-type. While the mean of *dSlo*^−^ photoreceptor output (blue) decayed slower than in the other mutant R1-R6s and undershot less.

Concurrently, the wild-type and mutant R1-R6 output ranges - measured as the standard deviation (Fig. 6*D*) of their response waveforms (Fig. 6*C*) at each second of NS - adapted with distinctive dynamics and speeds. *dSlo*^−^ R1-R6 outputs desensitized the slowest, slower than the wild-type, with their ranges compressing with different average time courses (T_dSlo-_ = 3.41 ± 3.28 s, n = 19 cells [22 recordings]; T_Wild-type_ = 1.47 ± 0.67, n = 7 cells [10 recordings]; mean ± SD) (Fig. 6*D*). Conversely, *dSK*^−^ and *dSK*^−^;;*dSlo*^−^ R1-R6 output ranges first compressed as rapidly as the wild-type (T_dSK-_ = 1.45 ± 0.66, n = 7 cells [7 recordings]; T_dSK-;;Slo-_ = 1.44 ± 0.32, n = 8 cells [9 recordings]), but then slowly begun to expand, reflecting their rather similar mean voltage dynamics (Fig. 6*B*). The adaptive range reduction occurred most severely in *dSK*^−^;;*dSlo*^−^ and *dSlo*^−^ R1-R6s, leaving their steady-state responses ~10% smaller than those of the wild-type.

These results highlight the complex role of Ca^2+^-activated K^+^ channels in regulating R1-R6 output in network adaptation. While the absence of dSlo channel slowed adaptation in *dSlo*^−^ R1-R6s, the *dSK*^−^ and the double-mutant *dSK*^−^;;*dSlo*^−^ R1-R6s adapted faster but showed overshooting dynamics. Consequently, as an overall sign of compromised gain control, the mutant R1-R6s reached their steady-state responsiveness 20-30 s later than the wild-type. Thus, each mutant R-LMC-R system adapted suboptimally, constrained to its own unique dynamics.

### dSK or dSlo Absence Leaves Information Sampling Intact

A R1-R6’s information transfer rate depends mostly on its photon-absorption rate changes, set by the number of individual sampling units (rhabdomeric microvilli) and the speed and refractoriness of their phototransduction reactions (Song et al., 2012; Juusola et al., 2017; Juusola and Song, 2017). In contrast, obeying the data processing theorem, any changes in membrane filtering affect signal and noise equally, and therefore cannot increase information (Shannon, 1948; Juusola and de Polavieja, 2003; Cover and Thomas, 2006). Accordingly, information transfer rates of mutant photoreceptors with normal phototransduction but without specific K^+^ channels, such as the slow delayed rectifier Shab (I_KS_) (Vähäsöyrinki et al., 2006), are broadly wild-type-like. But mutations that damage ion channels can destroy information. For example, *Sh* mutant R1-R6s’ “nonfunctional” *Shaker* (I_A_) K^+^ channels appears to truncate signal amplification while generating noise, reducing information flow (Niven et al., 2003). Critically, however, the R-LMC-R system has intrinsic potential to combat detrimental changes within its parts. A R1-R6’s impaired function can be compensated in part by extra light information (through gap-junctions and feedback synapses) from its neighbors, in which receptive fields face the same visual area (Shaw, 1984; Zheng et al., 2006; Wardill et al., 2012; Juusola et al., 2017).

Because *dSlo*^−^, *dSK*^−^ and *dSK*^−^;;*dSlo*^−^ mutant R1-R6s lack completely their functional channels (which thus should not generate extra noise) and have normal rhabdomere morphology and LIC dynamics (Fig. 5), theoretically, their somatic information transfer rates should be wild-type-like, or slightly lower; in case, their LMC feedback was compromised.

To test this hypothesis, we compared *dSlo*^−^, *dSK*^−^ and *dSK*^−^;;*dSlo*^−^ R1-R6s’ encoding performance to the wild-type control using the same recordings as above. In each case, the first 20-30 responses with the adapting trends were removed. The signal was taken as the average of the next 20 responses, which thus had settled to a relative steady-state, with its power spectrum calculated by Fourier transform. The corresponding noise power spectrum was estimated from the difference between each response and the signal (see Materials and Methods).

We found that the mutant R1-R6s’ signal-to-noise ratios (Fig. 6*E*) and information rates (Fig. 6*F*) were broadly wild-type-like; increasing in parallel with brightening light, as tested for dim, middle and bright NS. Thus, as hypothesized, after the initial ~20-30 s adaptation phase, the loss of dSK, dSlo or both channels affected only marginally a R1-R6’s encoding performance. These results highlight the R-LMC-R system’s robustness and compensatory ability to withstand internal damage.

### dSK or dSlo Absence Increases Synaptic Feedback

To work out in theory how synaptic feedback from the lamina interneurons should shape the wild-type R1-R6 output and how homeostatic feedback changes should shape mutant R1-R6 outputs, we next combined biophysical R1-R6 modelling with intracellular recordings.

Our biophysical R1-R6 model (Fig. 7*A*) incorporates 30,000 computational microvilli (Song et al., 2012), each of which implements full stochastic phototransduction reactions to transduce absorbed photons into QBs. Essentially, this model samples light information much like a real R1-R6 (Song et al., 2012; Song and Juusola, 2014; Juusola et al., 2017; Juusola and Song, 2017). Its QBs sum up realistic macroscopic LIC, with the best performance for naturalistic stimuli at 1-8 × 10^5^ photon absorptions/s (Song and Juusola, 2014; Juusola et al., 2017). LIC then charges a Hodgkin-Huxley-type photoreceptor membrane circuit (Figs 7*B* and 8) (Niven et al., 2003; Vähäsöyrinki et al., 2006; Song et al., 2012; Song and Juusola, 2014), generating output that approximates intracellular recordings to comparable light stimulation (Song et al., 2012; Song and Juusola, 2014; Juusola et al., 2017; Song and Juusola, 2017). Most differences in the simulated and recorded response waveforms would then be caused by the real R1-R6s’ synaptic feedback currents - input from LMCs (Zheng et al., 2006; Rivera-Alba et al., 2011; Dau et al., 2016), which the model lacks (Juusola et al., 2017). Moreover, given that the mutant R1-R6s’ phototransduction is wild-type-like and voltage-sensitive conductances either wild-type-like or only moderately reduced (Fig. 5), their voltage response differences should also mostly reflect synaptic feedback differences (Fig. 6).

**Figure 7.**
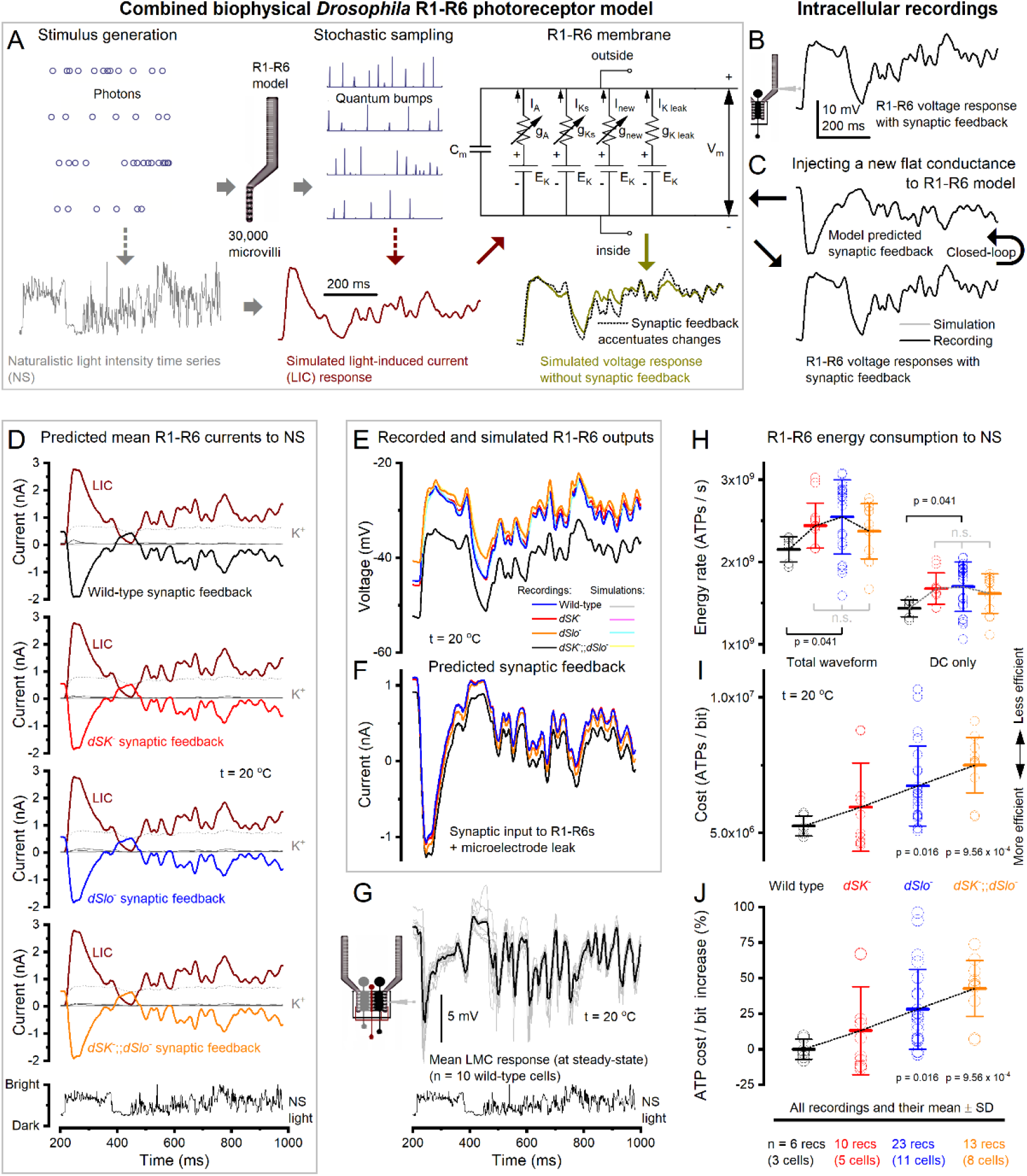
Predicted Synaptic Feedback and ATP Consumption of Wild-Type and Mutant R1-R6 Photoreceptors. ***A***, Biophysically realistic R1-R6 model has four modules: stimulus generation, stochastic photon sampling/quantum bump generation, bump integration and voltage-sensitive membrane. But it lacks the synaptic feedback from the lamina network (Juusola et al., 2017; Song and Juusola, 2017), which affect the real R1-R6 output. R1-R6 simulations (dark yellow trace) and recordings (dotted trace) to a repeated naturalistic light intensity time series (NS) were analyzed at relative steady-state adaptation (*cf*. Fig. 6*C*). ***B***, Characteristic recording waveform to bright NS (BG0). ***C***, Synaptic feedback to each recording was estimated computationally by linking it to the photoreceptor model, which had no free parameters. A new flat (zero) conductance, representing the synaptic input, was then injected to the model. This conductance waveform was shaped in a closed-loop until the model output (gray) matched the recorded output (black). ***D***, The fixed light-induced (dark red), K^+^ currents and the average predicted synaptic feedback and of wild-type and mutant R1-R6 recordings. ***E***, Together, these currents charged up their respective simulated R1-R6 voltage responses. The simulations (light colors) match the recordings (bright colors) near perfectly. ***F***, The average predicted synaptic feedback was unique to the mutant R1-R6s and showed stronger modulation on a higher mean (tonic excitatory background) than the wild-type (see also Fig. 10*F*). Testing the feedback means across all recordings: wild-type vs *dSK*^−,^ p = 0.041; *dSK*^−^ vs *dSlo*^−^, p = 0.033; *dSlo*^−^ vs *dSK*^−^;;*dSlo*^−^, p = 0.009; testing the mean feedback waveforms against each other, p < 2.274 × 10^−62^. ***G***, Separately recorded large monopolar cell (LMC) response waveforms to the same NS much resemble the predicted feedback waveforms (in F), suggesting that L2 and L4 cells, which form feedback synapses with R1-R6s (Meinertzhagen and O’Neil, 1991; Rivera-Alba et al., 2011), would contribute to R1-R6 output modulation (Zheng et al., 2006). ***H***, With these conductances included in each separate wild-type, *dSK^−^, dSlo*^−^ and *dSK-;;dSlo*^−^ R1-R6 models, the metabolic energy (ATP) consumption of each recording was calculated for its full waveform (left) (Song and Juusola, 2014) and DC voltage (right) (Laughlin et al., 1998), respectively. Notably, the original DC voltage method, which does not consider how the dynamic ion fluctuations add to the electrochemical pumping work, underestimates ATP consumption by 1/3 (33.2%; see Materials and Methods). ***I***, The cost of neural information, was calculated for each recording by dividing its information rate estimate with its full ATP consumption rate estimate. On average, the absence of dSK or dSlo or both increased the cost of neural information in a mutant R1-R6 by 13.1% (*dSK*^−^), 28.0% (*dSlo*^−^) or 42.7% (*dSK*^−^;;*dSlo*^−^).

Therefore, we could extrapolate the synaptic feedback current to each recorded R1-R6, whether wild-type or mutant, computationally (Fig. 7*C*) by using the same fixed LIC with their specific I_A_ and I_SK_ current dynamics (Figs 5 and 8). In these simulations, we first injected a new flat (zero) conductance, representing the missing synaptic input, to the full R1-R6 model. The software then shaped up this conductance waveform in a closed-loop until the model’s voltage response matched the recorded response for the same light stimulus. Thus, theoretically, the resulting (predicted) current should closely mimic the real synaptic feedback, which the tested R1-R6 would have received from the lamina network *in vivo*.

**Figure 8.**
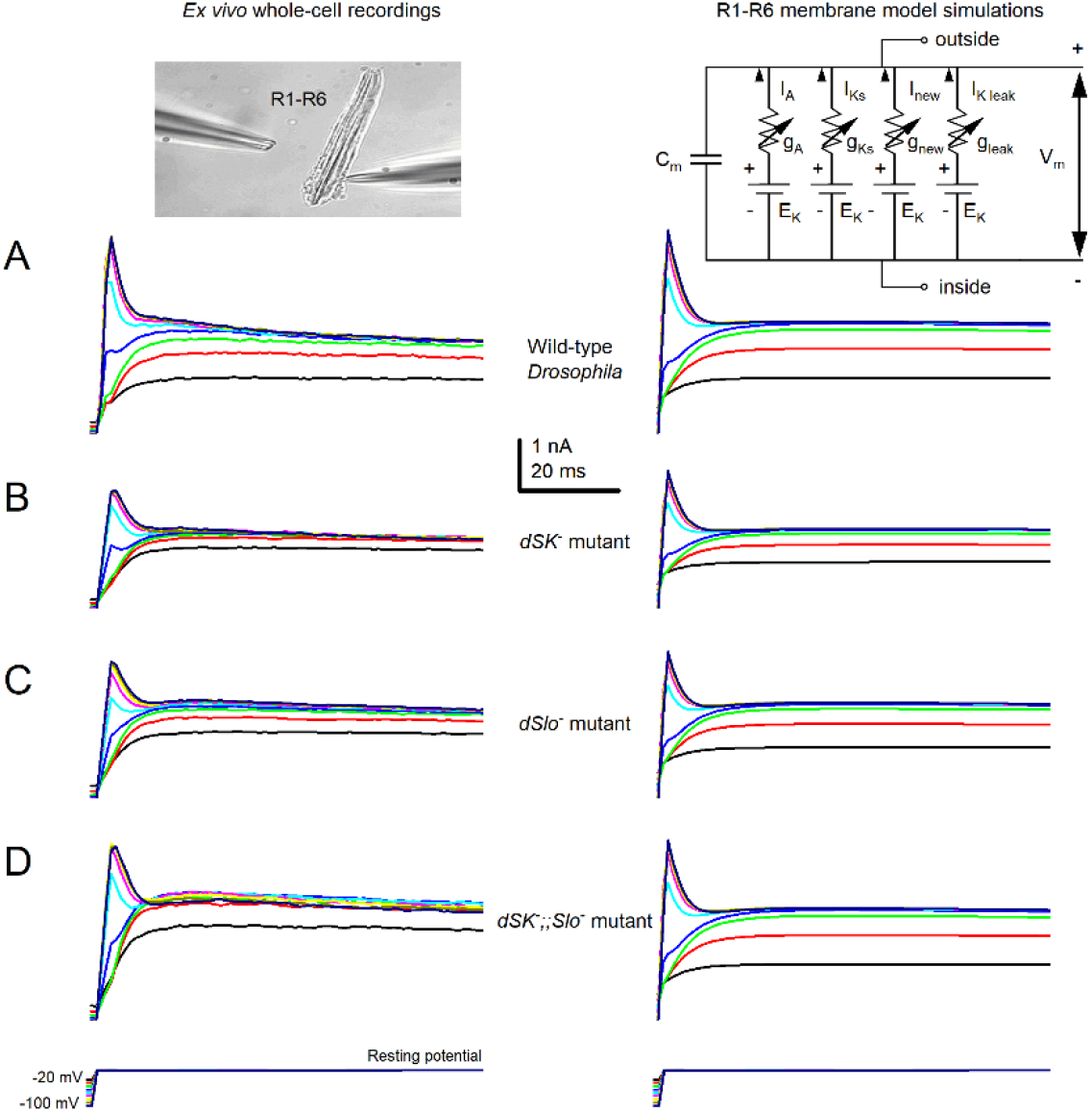
R1-R6 photoreceptors’ characteristic voltage-sensitive *Shaker* and *Shab* K^+^ current responses *ex vivo* and their HH-models to voltage commands, under whole-cell voltage-clamp conditions. ***A***, wild-type ***B***, *dSK*^−^ ***C***, *dSlo*^−^ ***D***, *dSK^−^;;dSlo*^−^

Fig. 7*D* shows the corresponding mean LIC and synaptic feedback estimates to repeated light stimulation for the tested wild-type and mutant photoreceptors, and the concurrent voltage-sensitive K^+^ currents and K^+^ leak estimates. In these simulations, whilst the LIC was the same (fixed; dark red traces) for every genotype, their synaptic feedback and K^+^ (dark green) currents balanced out differently to reproduce their respective *in vivo* voltage signals (Fig. 7*E*).

We found that in every simulation the predicted synaptic feedback to R1-R6s was excitatory, graded and phasic (Figs 7*D* and 7*F*). It rapidly increased (“switched-on”) during light decrements and decreased (“switched-off”) during light increments. This accentuated transient (phasic) light changes in photoreceptor output (Fig. 7*E*; *cf*. Fig. 7*A*). Moreover, the predicted synaptic excitatory load to R1-R6s (Fig. 7*F*) was unique for each mutant and the wild-type flies with the highest mean to *dSK*^−^ (red) and *dSlo*^−^ (blue) photoreceptors. Thus, the enhanced excitatory feedback conductance from the lamina interneurons is the most probable mechanistic explanation of why and how the mutant photoreceptors were more depolarized than their wild-type counterparts, both in darkness (*cf*. Fig. 4*D*) and during light stimulation (Fig. 7*E*).

Remarkably, these feedback dynamics (Fig. 7*F*), which were extrapolated using only photoreceptor data (Figs 7*A-C*), closely resembled postsynaptic intracellular LMC responses to the same light stimulus (Fig. 7*G*). This implied that L2, L4 and lamina intrinsic amacrine neurons (Lai), all of which receive inhibitory inputs from R1-R6 but form excitatory feedback synapses to R1-R6 (Kolodziejczyk et al., 2008; Raghu and Borst, 2011; Hu et al., 2015; Davis et al., 2018), could alone or together be the major source of this feedback. Thus, these new findings are consistent with our theory of how the R-LMC-R system, by dynamically balancing its inhibitory and excitatory synaptic loads, shapes the early neural representation of visual information (Zheng et al., 2006; Nikolaev et al., 2009; Zheng et al., 2009; Dau et al., 2016).

### dSK and dSlo Lower Neural Information Energy Cost

In response to LIC and synaptic feedback, ion channels open and close, regulating the ionic flow across the photoreceptor membrane. Meanwhile its ion cotransporters, exchangers and pumps uptake or expel ions to maintain ionic concentrations in- and outside. The work of the pumps in moving ions against their electrochemical gradients consumes ATP (Laughlin et al., 1998). For a R1-R6, a reasonable estimate of this consumption can be calculated from the ionic flow dynamics through its ion channels; details in Materials and Methods (see also: Song and Juusola, 2014).

Using our biophysical R1-R6 model, which now included the synaptic feedback, we calculated how much each recorded wild-type and mutant R1-R6 consumed metabolic energy (ATP molecules/s) to encode bright naturalistic light changes (Fig. 7*H*, left). We discovered that because their enhanced synaptic feedback held *dSK*^−^, *dSlo*^−^ and *dSK*^−^;;*dSlo*^−^ R1-R6s at higher operating voltages, where signaling is more expensive, they consumed on average 13.3%, 18.3% and 10.2% more ATP than the wild-type, respectively.

We also estimated each tested R1-R6’s ATP consumption by using the method of balancing out the ionic currents for its light-induced mean (flat) depolarization level, or DC (Laughlin et al., 1998). This produced a metric, which followed quite a similar trend (Fig. 7*H*, right). But because it discarded how much the dynamic ion fluctuations increase the work to maintain transmembrane ionic concentration, it underestimated the total ATP consumption by ~1/3.

Next, using the full biophysical models (Fig. 9), we calculated how the mutant R1-R6s’ homeostatically reduced *Shaker* and *Shab* K^+^ conductances (Figs 5*F-H*) affect their neural information costs. We fixed the *Shaker* and *Shab* conductance dynamics of the *dSK*^−^, *dSlo*^−^ and *dSK*^−^;;*dSlo*^−^ R1-R6 models to match typical wild-type R1-R6 VC-recordings (Fig. 8*A*). This increased the mutant photoreceptors’ energy consumption, but only slightly (*cf*. Figs 9*H* to 7*H*). Hence, the observed homeostatic 19-36% *Shaker* and *Shab* current reduction in *dSK*^−^ and *dSlo*^−^ R1-R6s (Figs 5*F* and 5*H*) made evolutionary sense, as it cut both their hyperpolarizing drive, which therefore would require less excitatory synaptic feedback to depolarize the cells, and neural information costs. But this saving was small, only 4.5-6.2%. And somewhat unexpectedly, its homeostatic effect, in fact, increased the *dSK*^−^ and *dSlo*^−^ R1-R6s’ synaptic feedback overload slightly in respect to *dSK*^−^;;*dSlo*^−^ R1-R6s, which had wild-type-like *Shaker* and *Shab* conductance dynamics (Fig. 7*F*).

**Figure 9.**
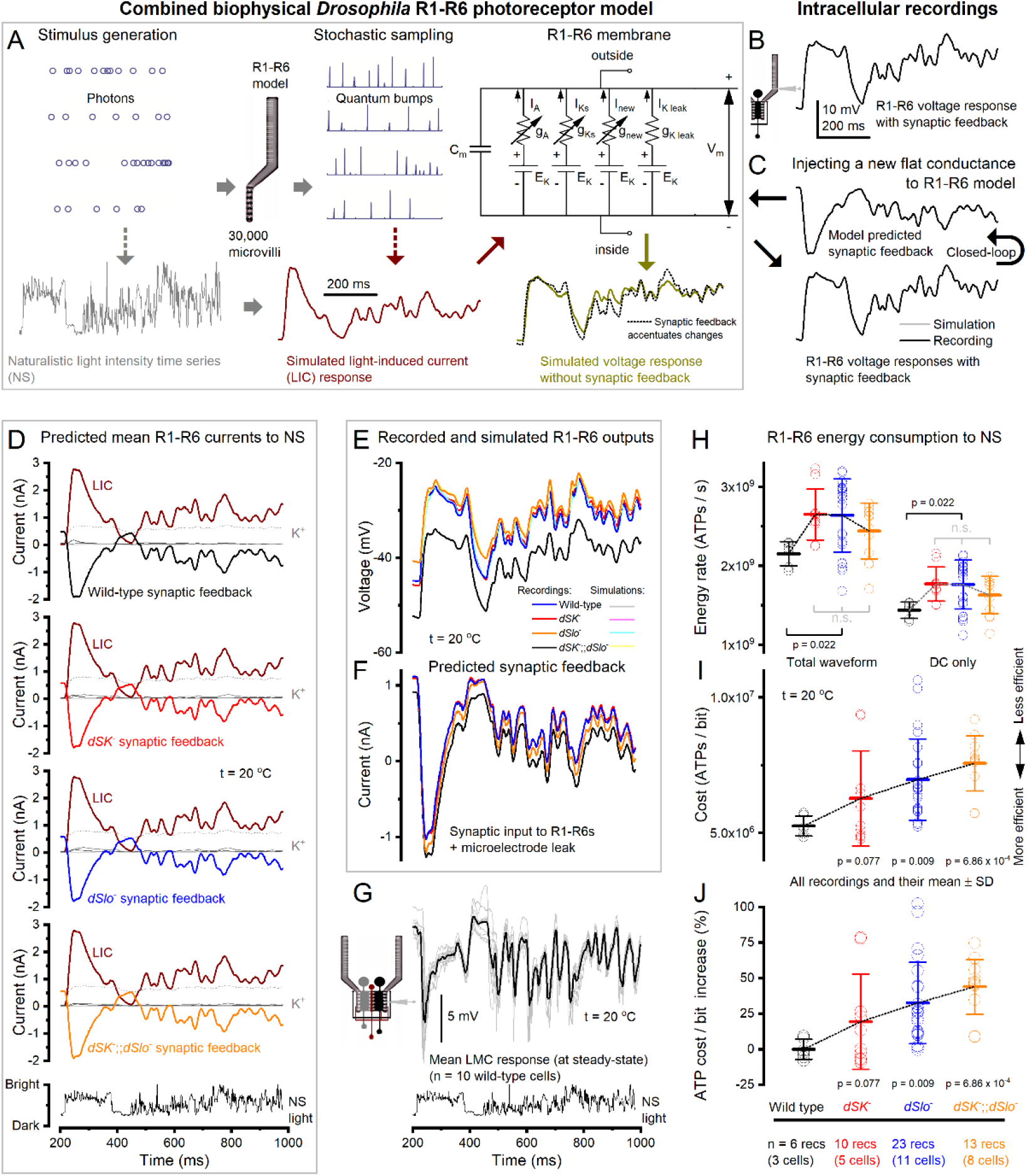
Examples of the wild-type (black), *dSK*^−^ (red), *dSlo*^−^ (blue) and *dSK*^−^;;*dSlo*^−^ (orange data) photoreceptor models’ *Shaker* (I_A_), *Shab* (I_Ks_), K^+^-leak, LIC and synaptic feedback currents to naturalistic light stimulation, when using the same wild-type *Shaker* and *Shab* conductance dynamics in all the models; *cf*. Figs 7 and 10. ***A***, In these simulations, the while-type, *dSK*^−^, *dSlo*^−^ and *dSK*^−^;;*dSlo*^−^ R1-R6 models had identical light induced current (LIC) and voltage-sensitive membrane conductances (the models used the wild-type *Shaker* and *Shab* dynamics as in Fig. 8*A*; *cf*. Fig. 7*A*). ***B***, Characteristic recording waveform to bright NS (BG0). ***C***, Again, synaptic feedback was computed through the R1-R6 model, which had no free parameters, in a closed-loop until the model output (gray) matched the recorded output (black). ***D***, The fixed light-induced (dark red), K^+^ currents and the average predicted synaptic feedback of wild-type and mutant R1-R6 recordings. ***E***, Together, these currents charged up their respective simulated R1-R6 voltage responses. The simulations (light colors) matched the recordings (bright colors). ***F***, Again, the average predicted synaptic feedbacks to the mutant R1-R6s were stronger, having higher means (tonic excitatory background) than the wild-type. ***G***, The recorded large monopolar cell (LMC) response waveforms to the same NS resembled the predicted feedback waveforms (in F). ***H***, The ATP consumption of these mutant R1-R6 models was 3.5-8.6% higher than in those models, which had their recorded (smaller) *Shaker* and *Shab* conductances (*cf*. Figs 5*F-H* and 7*H*). ***I***, The cost of neural information, was calculated for each recording by dividing its information rate estimate with its full ATP consumption rate estimate. On average, the absence of dSK or dSlo or both increased the cost of information in a mutant R1-R6 by 19.3 ± 33.4% (*dSK*^−^), 32.6 ± 28.6% (*dSlo*^−^) or 43.9 ± 19.3% (*dSK*^−^;;*dSlo*^−^). Thus, 19-36% homeostatic reductions in *Shaker* and *Shab* currents in *dSK*^−^ and *dSlo*^−^ R1-R6 photoreceptors caused only 4.5% and 6.2% savings in their ATP consumption per each bit of transmitted information (*cf*. Figs 7*I* and 7*J*).

Furthermore, simulations about other possible homeostatic changes (Fig. 10) indicated that by increasing leak and voltage-sensitive K^+^ conductances, or adding an extra Cl^−^-leak, in the R1-R6 membrane would strengthen and accentuate synaptic feedback (Fig. 10*F*), and by that increase both the wild-type R1-R6s’ ATP consumption (Fig. 10*H*; now by 23.2%) and the mutant photoreceptors’ neural information costs in respect to the wild-type (Figs 10*I* and 10*J*), now by 22.3% (*dSK*^−^), 37.0% (*dSlo*^−^) or 57.6% (*dSK*^−^;;*dSlo*^−^). Therefore, as energy wasting reduces fitness, the earlier proposed leak-conductance overexpression alone (Niven et al., 2003; Vähäsöyrinki et al., 2006) seems an unlikely homeostatic strategy here.

**Figure 10.**
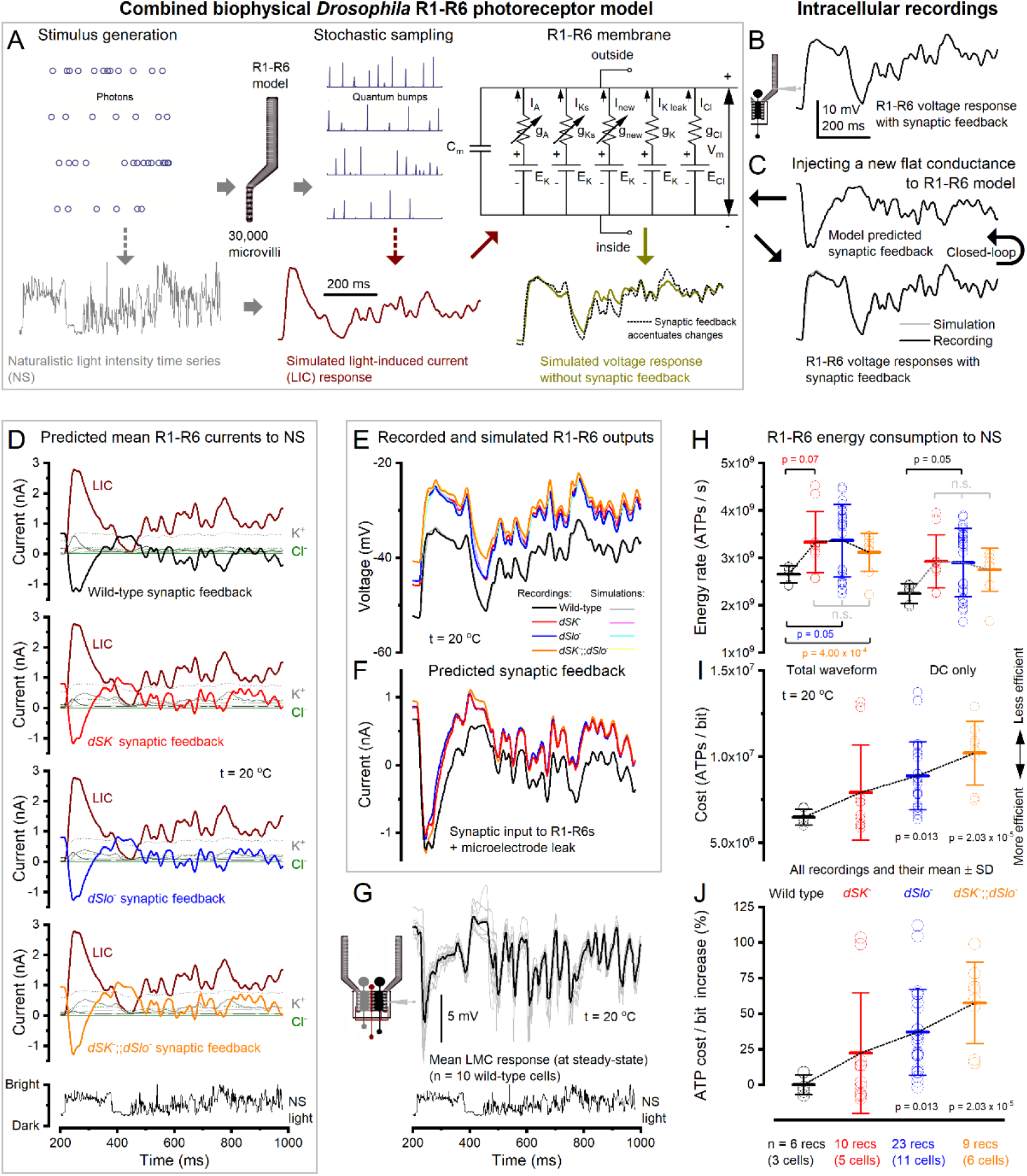
Examples of the wild-type (black), *dSK*^−^ (red), *dSlo*^−^ (blue) and *dSK*^−^;;*dSlo*^−^ (orange data) photoreceptor models’ *Shaker, Shab*, K^+^-leak, Cl^−^-leak LIC and synaptic feedback currents to naturalistic light stimulation, when using larger wild-type *Shaker* and *Shab* conductance dynamics (as in Table 2) in all the models; *cf*. Figs 7 and 9. ***A***, In these simulations, we further added a Cl^−^-leak and Cl^−^ conductance (here combined to g_Cl_^−^ to simply the HH-diagram) in the R1-R6 photoreceptor membrane model and balanced these by increasing voltage-sensitive K^+^ conductances (Materials and Methods: speculative photoreceptor membrane model), and again the synaptic feedback (Juusola et al., 2017; Song and Juusola, 2017) was computed in a closed-loop until the simulations matched the recordings (*cf*. Fig. 6*C*). ***B***, Characteristic recording waveform to bright NS (BG0). ***C***, Synaptic feedback to each recording was estimated computationally by linking it to the photoreceptor model, which had no free parameters. ***D***, The fixed light-induced (dark red), K^+^ and Cl^−^ currents and the average predicted synaptic feedback and of wild-type and mutant R1-R6 recordings. ***E***, These currents charged up their respective simulated R1-R6 voltage responses (light colors), which matched the actual recordings (bright colors). ***F***, Similar to the other simulations (*cf*. Figs 7*E* and 9*E*), the predicted synaptic feedback to the mutant R1-R6s was larger (carrying bigger modulation) with a higher mean (tonic excitatory background) than the wild-type. However, the modulation in these simulations was even more transient. ***G***, Separately recorded large monopolar cell (LMC) response waveforms to the same NS much resemble the predicted feedback waveforms (in F). ***H***, Energy (ATP) consumption of each recording was calculated for its full waveform (left) (Song and Juusola, 2014) and DC voltage (right) (Laughlin et al., 1998), respectively. Notably, the added extra Cl^−^-leak and Cl^−^ conductance (with rebalanced K^+^ conductances) increased the photoreceptors’ energy usage by ~26.3% (*cf*. Fig. 7*H*, left: from 2.10 × 10^9^ ATP/s to 2.65 × 10^9^ ATP/s,). Whilst the original DC voltage method (right), which does not consider how the dynamic ion fluctuations add to the electrochemical pumping work, now underestimated ATP consumption by about 15%. ***I***, The cost of neural information, was calculated for each recording by dividing its information rate estimate with its full ATP consumption rate estimate. Here, homeostatic increase in K^+^ and Cl^−^ leak conductances (to compensate the loss of Ca^2+^-activated K^+^ channels) increased the cost of information in a mutant R1-R6 by 22.2 ± 42.3% (*dSK*^−^), 37.0 ± 30.3% (*dSlo*^−^) or 57.6 ± 28.8% (*dSK*^−^;;*dSlo*^−^), in respect to the comparable wild-type model.

These results establish the extra energy, which a mutant R1-R6 must spend to function without Ca^2+^-activated K^+^ channels, as a major cost for homeostatic compensation of neural information (Fig. 7*I*). To maintain similar information rates (Fig. 6*F*), an average mutant R1-R6 consumed at least 13.1% (*dSK*^−^; p = 0.114), 28.0% (*dSlo*^−^; p = 0.016) or 42.7% (*dSK*^−^;;*dSlo*^−^; p = 9.56 × 10^−4^) more ATP for each transmitted bit than its wild-type counterpart (Fig. 7*J*). Notably, these costs would only increase further if homeostatic compensation of the missing dSK and dSlo channels further entailed overexpression of additional K^+^ or Cl^−^ conductances or leaks (Figs 9 and 10). Thus, in *Drosophila* photoreceptors, Ca^2+^-activated K^+^ channels reduce the energy cost of neural information.

### dSlo and dSK Co-Regulate Feedforward Transmission to LMCs

Thus far, we have provided experimental and theoretical evidence that both BK (dSlo) or dSK channel deletions enhance synaptic feedback from the lamina interneurons to R1-R6s (Figs 3–10). But these results still leave open the corresponding changes in the post-synaptic LMC output, which initiates the motion vision pathways to the fly brain (Joesch et al., 2010; Wardill et al., 2012). To test how dSK and dSlo deletions affect such feedforward transmission directly, we recorded intracellular voltage responses of dark-adapted LMCs in the mutant and wild-type laminae to brightening light flashes, which covered a 4-log intensity range (Fig. 11*A*).

**Figure 11.**
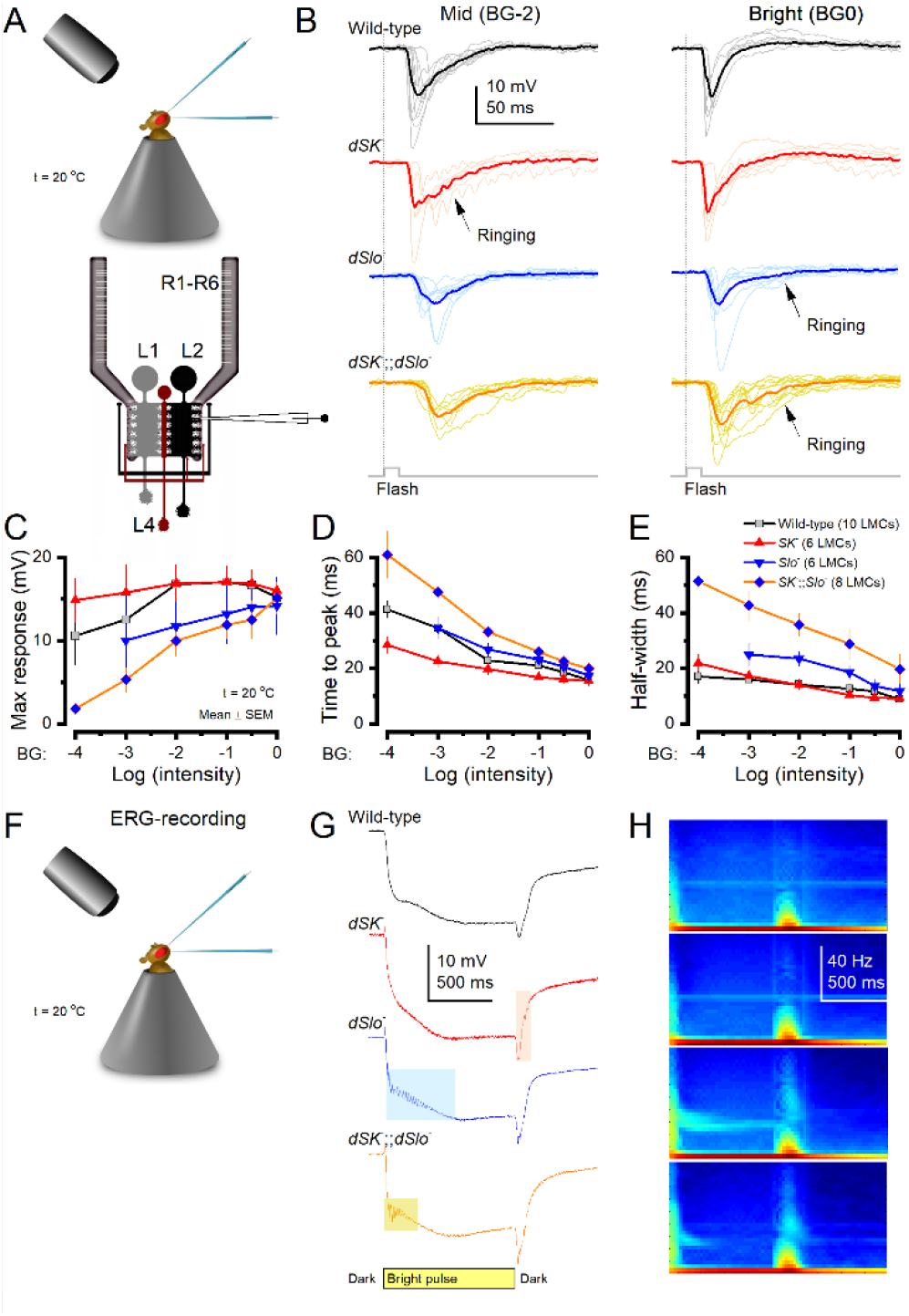
Large Monopolar Cell (LMC) Output in the Wild-Type and Mutant Flies Differ Consistently. ***A***, Intracellular LMC recordings were carried out from *in vivo* (above). The schematic (below) highlights synaptic feedbacks from L2/AC and L4 to photoreceptors terminals. ***B***, Voltage responses of the wild-type and mutant LMCs to Bright and Middle intensity flashes. ***C***, *dSlo*^−^ and *dSK*^−^;;*dSlo*^−^ LMCs generated smaller responses to the dim and middle intensity flashes than the wild-type (*dSlo*^−^: p < 0.01, BG0-4, n = 8; *dSK*^−^;;*dSlo*^−^: p < 0.03, BG0-4, n = 5). ***D***, *dSK*^−^ LMC responses peaked the fastest, whereas *dSlo*^−^ and *dSK*^−^;;*dSlo*^−^ LMC responses were slower than the wild-type (p < 0.03) over the tested intensity range. ***E***, *dSK*^−^ LMC responses lasted as long as the wild-type responses, while *dSlo*^−^ and *dSK*^−^;;*dSlo*^−^ LMCs took much longer to repolarize than wild-type (*dSlo*^−^, p < 0.04, BG2-3; *dSK*^−^;;*dSlo*^−^, p < 0.04, BG0-3). ***F***, *In vivo* electroretinograms (ERG), depicting global light-induced eye activity, were recorded from the corneal surfaces of intact *Drosophila*. ***G***, *dSK*^−^, *dSlo*^−^ and *dSK*^−^;;*dSlo*^−^ ERGs often showed characteristic oscillations after the light on- and off-transients, consistent with the corresponding intracellularly recorded light-induced LMC oscillations (B). ***H***, Dynamic spectra of the ERGs (G) reveal the frequency-dependency and duration of the oscillations. ***C-E***: Mean ± SEM, two-tailed t-test.

Expectedly, light rapidly hyperpolarized LMCs and darkness depolarized them (Fig. 11*B*) (Zettler and Järvilehto, 1973; Juusola et al., 1995a; Zheng et al., 2006), driven by the photoreceptors’ inhibitory transmitter, histamine (Hardie, 1989; Dau et al., 2016). Yet, these dynamics varied somewhat systematically between the genotypes, with the mutant LMCs often showing oscillating responses (ringing) around specific frequencies. L1 (on-pathway) and L2 (off-pathway) responses are thought to be largely similar at the dendritic (lamina) level (Hardie and Weckström, 1990; Uusitalo et al., 1995a; Nikolaev et al., 2009) (*cf*. Fig. 7*G*), with their medulla terminals’ light-on and -off preference (Joesch et al., 2010; Freifeld et al., 2013) most likely arise through specific medulla circuit processes. Therefore, with most penetrations likely from L1 and L2, which are the largest LMCs, our recordings should mostly depict mutation-induced variations and less LMC-type-dependent differences.

*dSK*^−^ LMC output was consistently the most transient, even to dim flashes (Figs 11*B-E*), showing accelerated (most “light-adapted”) dynamics with the fastest time-to-peak values (Fig. 11*D*). By and large, the size (Fig. 11*C*) and half-width (Fig. 11*E*) of these responses were wild-type-like, but, unlike the wild-type, they often showed rapid oscillation bursts to dim flashes (see also Abou Tayoun et al., 2011).

In contrast, both *dSlo*^−^ and *dSK*^−^;;*dSlo*^−^ LMC responses to dimmer flash intensities were on average smaller than those of the wild-type and *dSK*^−^ LMCs (Figs 11*B* and 11*C*). But as their amplitudes increased with light intensity, the brightest flashes evoked about the same size responses from all the genotypes (Figs 11*B* and 11*C*). Therefore, during dim (but not bright) stimulation, the excitatory feedback from L2 and L4 cells to R1-R6s (Zheng et al., 2006), if directly following the recorded *dSlo*^−^ and *dSK*^−^;;*dSlo*^−^ LMC responses, could be driven by smaller *dynamic* modulation (Fig. 11*C*) on a larger *static* load (as their mean LMC responses would thus also be more depolarized). This would reduce R1-R6 membrane impedance and, presumably, synaptic gain in R1-R6 output; consistent with the smaller *dSlo*^−^ and *dSK*^−^;;*dSlo*^−^ R1-R6 responses to dim naturalistic light stimulation (Fig. 6*C*). Furthermore, *dSK*^−^;;*dSlo*^−^ LMC response dynamics were also slower and less tightly time-locked (Fig. 11*D*); often ringing sluggishly (Fig. 11*B*), prolonging the response half-width (Fig. 11*E*) and peaking later than the other corresponding LMC responses (Fig. 11*D*). Such desynchrony would add noise in the synaptic feedback, and may have contributed to the slightly lower signal-to-noise ratios and information transfer rates of *dSK*^−^;;*dSlo*^−^ R1-R6s (Fig. 6*E*).

Thus, deletion of dSK, dSlo or both led to suboptimal network adaptation in the R-LMC system, seen as accelerated or decelerated LMC responses and mutation-specific oscillations. Crucially, these oscillations, with their characteristic frequencies, were also regularly observed in the mutant eyes’ global electrical activity (electroretinograms, ERGs) (Figs 11*F-H*), supporting the intracellular results. Nevertheless, the observed differences cannot be directly attributed for missing dSK, dSlo or both in the mutant LMCs (cf. Abou Tayoun et al., 2011). The respective functional channels (dSK in *dSlo-* mutants, dSlo in *dSK-* mutants, and both channels in wild-type) could act remotely in the circuit, or their LMC response dynamics could result from combinatorial effects on both R1-R6s and LMCs.

### Mutants’ Optomotor Responses Reflect Early Vision Defects

To test whether the mutation-specific network adaptations influence visual perception, we measured the flies’ optomotor behavior in a classic flight simulator system (Fig. 12*A*). Notably here, *dSK*^−^, *dSlo*^−^ and *dSK*^−^;;*dSlo*^−^ mutants lack their respective channel activity throughout their brains, and thus are likely to have perceptual deficits beyond their distorted LMC inputs - and in case of *dSlo*^−^, reduced health/motility (see Materials and Methods). But it is the LMC input, which sets their absolute motion vision limit (Rister et al., 2007; Joesch et al., 2010; Wardill et al., 2012). So, whilst any observed phenotype is convolving the channel contributions in some complex, unknown way across cell types, LMC input to the mutant brain is still its motion vision bottleneck, driving optomotor behaviors in a closed loop. Therefore, as a mutant’s optomotor behavior cannot be better than, and must ultimately reflect, its LMC input, it is informative to compare their respective optomotor response to their LMC input at different stimulus conditions to determine the generic behavioral differences across the different phenotypes. Moreover, these comparisons tell us further each mutant phenotype’s capacity to compensate its specific mutation effects.

**Figure 12.**
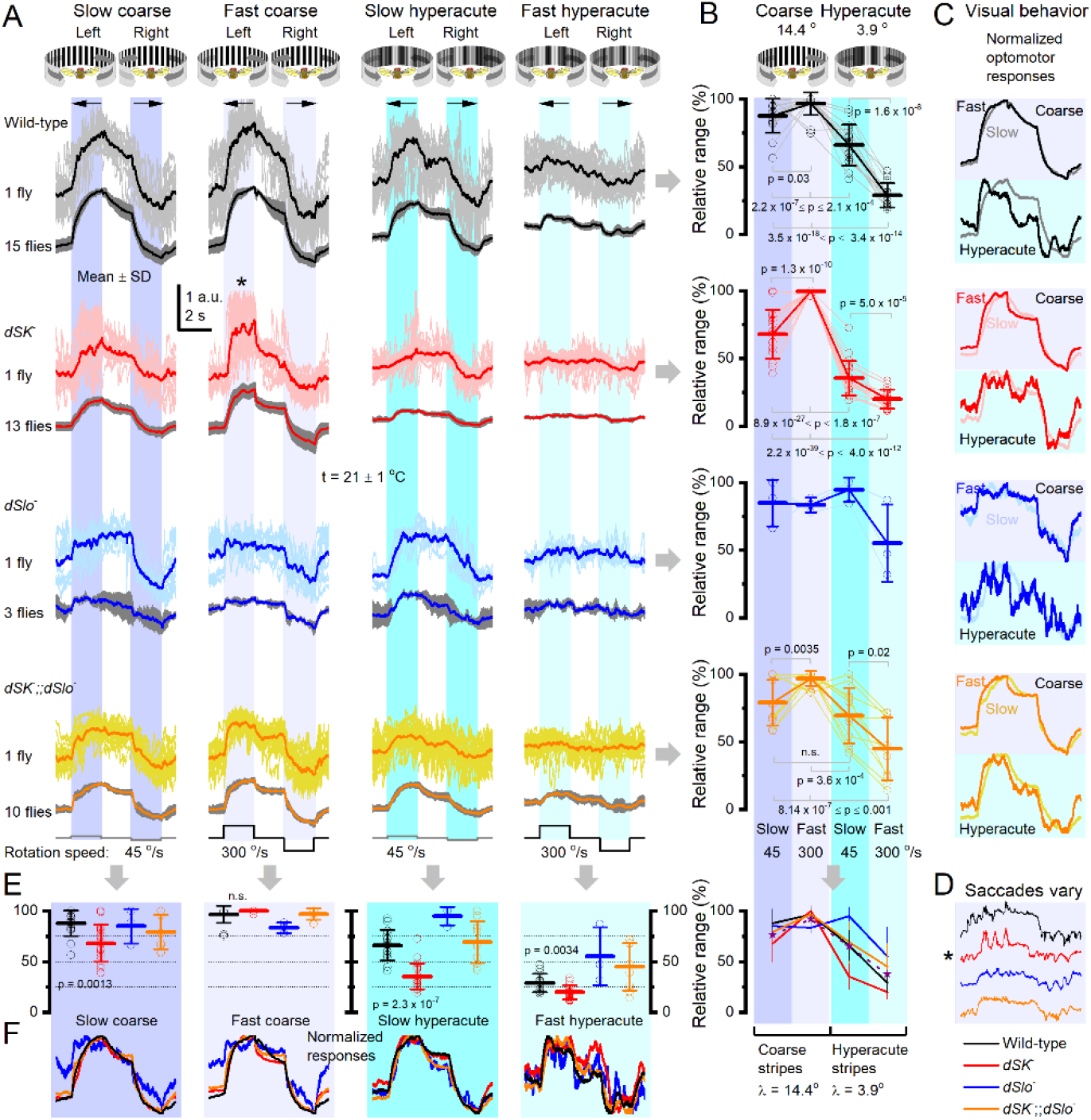
Wild-Type Flies and Ca^2+^-Activated K^+^ Channel Mutants Differ in Their Sensitivity to Track Different Field Rotations. ***A***, Optomotor responses to slow (45 °/s) and fast (300 °/s) left/right rotating fields of either coarse (14.4°) or finegrained (hyperacute: 3.9°) black-and-white vertical stripes (100% contrast), as seen by the flies. Above: mean (thick line) and 19-32 responses (thin lines) of the same fly to the four stimuli. Below: population means of many flies of the same genotype. Each fly was tested with these four stimuli to work out its relative optomotor sensitivity. In the experiments, the stimulus presentation order was actively varied to reduce novelty, flight fatigue or adaptation induced bias. ***B***, The relative output (%) range (max-min) of each tested fly, as scaled by its maximum optomotor response to the four stimuli. Wild-type, *dSK*^−^ and *dSK*^−^;;*dSlo*^−^ flies typically responded the strongest to the fast coarse field rotation, whilst *dSlo*^−^ flies were most sensitive to slow hyperacute field rotations. ***C***, Normalized responses of each genotype to each test stimuli. ***D***, The relative output range (max-min) of the different genotype of each test stimuli.. ***E***, Normalized responses of the different genotypes, compared to each other, for each test stimuli.

The tethered wild-type and mutant flies generated yaw torque by attempting to follow left and right rotating panoramic scenes, which showed either coarse (14.4°) or fine-grained (3.9°) vertical black-and-white stripe patterns, facing the flies. The resulting optomotor response waveforms and sizes were used to quantify how well individual flies and their respective populations (genotypes) saw these scenes rotating either slowly (45 °/s) or fast (300 °/s). Note that although the average inter-ommatidial angle (the eyes’ optical limit) is 4.5° (Gonzalez-Bellido et al., 2011), photomechanical photoreceptor microsaccades enable *Drosophila* to see much finer (hyperacute) details (Juusola et al., 2017).

We found that flies of each genotype could follow these stimuli (Fig. 12*A*), indicating that their visual systems represented and motor systems reacted to the opposing (left and right) image motion appropriately. However, the relative optomotor response sizes (Fig. 12*B*) and waveforms (Fig. 12*C*) showed genotype-specific sensitivities, or stimulus preferences, which were both repeatable and independent of the stimulus presentation order. Thus, these response differences could not be caused by stimulus salience, neural habituation or flight muscle fatigue.

Wild-type flies preferred, on average, the fast coarse stripe field rotations (Fig. 12*B*, black; 96.6 ± 8.5% maximum response, mean ± SD, n = 15 flies) over the slow coarse (87.8 ± 12.6%) and slow hyperacute (66.1 ± 15.2%) stimuli, but only just. Even their responses to fast hyperacute rotations were substantial (28.9 ± 9.0%), consistent with *Drosophila’s* high visual acuity even at saccadic speeds (>200 °/s) (Juusola et al., 2017). Such an all-round optomotor performance over a broad motion stimulus range is consistent with the idea that the optomotor behavior scales with the sensory input strength and dynamics from the eyes (Wardill et al., 2012). Thus, the adaptive signal scaling in the early visual system, seen as amplitude and time-normalized LMC contrast responses to different stimulus intensity and speed conditions (Zheng et al., 2006; Zheng et al., 2009), would be a prerequisite for consistent perception (optomotor behavior) over a broad image motion range.

In contrast, *dSK*^−^ mutants responded far more strongly to the fast coarse rotating field (Fig. 12*B*, red; 99.8 ± 7.6% maximum response, n = 13 flies) than the other stimuli (19.9-68.0%), with their slow and fast hyperacute field rotation responses being significantly weaker than those of the other genotypes (Fig. 12*E*). Interestingly and distinctively, the *dSK*^−^ responses were further dominated by large and fast body saccades (* in Fig. 12*A*), which appeared at seemingly regular intervals from the stimulus onset onwards and could make >50% of their total amplitude (Fig. 12*D*). Thus, the accelerated *dSK*^−^ photoreceptor and LMC dynamics (*cf*. Figs 3*C* and 11*C*), and tendency to oscillate, seem preserved in the *dSK*^−^ visual system, with these motion perception distortions possibly compelling their “spiky” optomotor responses.

The optomotor behavior of *dSlo*^−^ mutants showed similarly suggestive correlations to their R-LMC-R network adaptation dynamics. These flies, which boast slightly decelerated photoreceptor (Fig. 3*E*) and LMC (Fig. 11*D*) dynamics, preferred slow field rotations, and, surprisingly, were most sensitive to the slow hyperacute stimulus (Fig. 12*B*, blue; 94.8 ± 9.0%, n = 3 flies). Although *dSlo*^−^ mutants, in absolute terms, generated the weakest flight simulator torque responses of the tested genotypes, the mutants that flew did so over the whole experiments, making these stimulus preferences genuine.

Finally, the sensitivity of *dSK*^−^;;*dSlo*^−^ mutant responses (Fig. 12*B*, orange) followed the average of *dSK*^−^ and *dSlo*^−^ mutants’ optomotor responses (Fig. 12*B*, purple dotted line) more closely than the mean wild-type responses (black). In particular, their responses were relatively more sensitive to hyperacute stimuli than the corresponding wild-type responses (Fig. 12*E*) but rose and decayed slower (Fig. 12*F*, arrows), consistent with *dSK*^−^;;*dSlo* having slower LMC dynamics (Figs 11*D-E*). Thus, suggestively, their optomotor dynamics differences reflected more differences in early visual network adaptations rather than in other systems, such as the sensorimotor.

## Discussion

Our results indicate that dSlo (BK) and dSK (SK) reduce excitability and energy (ATP) consumption while increasing adaptability and dynamic range for transmitting neural information at the lamina network, ultimately stabilizing visual perception in changing light conditions. Here, single- and double-mutant photoreceptors showed either accelerated or decelerated responses and more depolarized resting potentials during steady-state adaptation. Such changes likely emerged from suboptimal homeostatic rebalancing of synaptic feed-forward and feedback signaling between photoreceptor axon terminals and the rest of the lamina network. Notably, this network compensation was unique for each mutation, resulting in distinctive adaptive regimes; with their respective LMCs showing oscillating accelerated or decelerated responses with reduced output ranges. These altered LMC response dynamics, and thus the flow of visual information, most probably distorted the mutants’ rotating scene perception, and their optomotor responses, in relation to the wild-type.

### Homeostatic Compensation Shapes both Electrical Responses and Synaptic Release

Because of the continuous bidirectional adapting interactions between photoreceptors and different lamina interneurons, the altered LMC responses cannot be explained simply by the absence of dSK and dSlo channels in the LMCs. In particular, both *dSlo* and *dSK* genes are well expressed in all LMCs (L1-L5) and photoreceptors (R1-R8) (Davis et al., 2018), underlying the interdependence of R1-R6 and LMC response dynamics and the need for systems level analyses to untangle them. Moreover, different neurons’ expression levels in the lamina terminals could vary dynamically, be tuned by circadian clock (Agrawal et al., 2017) or influenced (up- or down-regulated) by the Gal4-lines used to identify the cells (e.g. see I^A^ and I_Ks_-currents in Fig. 1). Nevertheless, irrespective whether these processes happen or not, homeostatic changes in the mutant R-LMC-R systems must involve both R1-R6s’ and LMCs’ electrical response waveforms and their synaptic release machineries. For example, in the *dSK*^−^ R-LMC-R system, the homeostatically rebalanced synaptic feedforward-feedback interactions (Fig. 7) and reduced R1-R6 Shaker and Shab K^+^ conductances (Fig. 5) alone would make their electrical response waveforms (Fig. 11*B*) different from the wild-type; as seen in *dSK*^−^ LMC waveforms peaking faster (Fig. 11D) and often oscillating to dim light.

### Ca^2+^-activated K^+^ Channels Reduce Costs of Adaptation and Increase Its Range

Adaptability is critical for animal fitness. In sampling and transmission of sensory signals, it reduces communication errors, such as noise and saturation, by continuously adjusting new responses by the memories of the past stimuli (Song et al., 2012; Juusola and Song, 2017). To ensure reliable perception of visual objects in changing conditions, retinal adaptation exploits visual world similarities and differences (van Hateren, 1992a; Song and Juusola, 2014) through characteristic visual behaviors (Schilstra and Hateren, 1999; Blaj and van Hateren, 2004; Juusola et al., 2017) and employs costly codes (de Polavieja, 2002) through multiple layers of feedbacks. This gives emergence for *homeostatic* network gain regulation, in which photoreceptor adaptation is mediated both by intrinsic (Juusola and Hardie, 2001b; Vähäsöyrinki et al., 2006; Song et al., 2012; Hardie and Juusola, 2015) and synaptic feedbacks (Zheng et al., 2006; Zheng et al., 2009). Here, the absence of dSK, dSlo or both channels left the phototransduction cascade essentially intact but reduced the intrinsic photoreceptor *Shaker* and *Shab* conductances, which should have made voltage responses larger and slower. Yet, *in vivo* recordings refuted these predictions, showing instead distinctive mutation-specific dynamics. Therefore, the observed defects in photoreceptor adaptability - including response fluctuations and altered dynamic ranges - seem mostly attributable to the R-LMC-R system’s suboptimally balanced synaptic feedforward inhibition and feedback excitation; reflecting homeostatic compensation at the network level. The resulting excitatory feedback overload also provided a plausible explanation why the mutant photoreceptors’ resting potentials and response speeds differed from the wild-type (Zheng et al., 2006; Abou Tayoun et al., 2011).

The primary effects of mutations can be difficult to separate from the secondary effects of homeostatic compensation (Marder and Goaillard, 2006). Nonetheless, the overall consistency of our findings suggest that many differences in *in vivo* response properties of the mutants’ R1-R6s and LMCs result from homeostatic gain regulation, whereupon differently balanced synaptic excitatory and inhibitory loads in the lamina network generate unique adaptive dynamics (encoding regimes); see also (Abbott and Lemasson, 1993; Lemasson et al., 1993). In the double-mutant, the most depolarized photoreceptors (Fig. 4*D*) and the slowest LMC output (Figs 11*D-E*) imply that the network gain was particularly challenging to regulate, providing the most compromised adaptability and response range (Fig. 12). In the single-mutants, adaptability of early vision was better compensated by enhanced network excitation, as seen by more wild-type-like LMC response dynamics (Figs 11*C-E*). But this still came with the cost of increased ATP consumption (Figs 7*H-I*). Moreover, in each case, the dSK and/or dSlo channel deletions affected optomotor behavior (Fig. 12), suggesting that the mutants’ distinct LMC output dynamics distorted their motion perception; alike what we have previously shown to occur with different color channel mutants (Wardill et al., 2012). Here, *dSK*^−^ mutants’ accelerated LMC responses (Figs 11*B-C*) presumably drove their fast hyper-saccadic optomotor responses (Figs 12*A-D*), while *dSlo*^−^ mutants’ decelerated LMC responses (Figs 11*B-C*) most probably sensitized their vision to slow scene rotations (Figs 12*A-D*).

We have shown how Ca^2+^-activated K^+^ channels serve local and global neural communication, improving economics and adaptability. Locally, they help to reduce calcium load and repolarize membrane potentials in synaptic terminals. Globally, they reduce the overall network excitability and the cost of transmitting information, while increasing the range of neural adaptation and reliable perception.

### Genetic control limitations

Finally, our results showed that the standard genetic rescue controls themselves - by using Gal4-lines and RNAi - can affect cellular form and function, causing larger neural response variability than what is observed in the tested phenotypes (Fig. 1). Thus, we could not use such controls to make *reductionist* conclusions about information processing at the network level; when analyzing neural response and homeostatic compensation dynamics in fine detail. These results highlight the importance of carefully testing the viability and usefulness of the planned control methods, both at the cellular and systems (network) level, so that the scientific rationale and reliability of the study becomes defined *constructionistically*, within the experimental/methodological limits.

## Author Contributions

This research was initiated by M.J., P.D. and R.C.H. Genetics: A.A.T., P.D. and F.B. Electrophysiology: X.L., A.D., D.R., A.N., L.Z., M.B., B.C., R.C.H. and M.J. Electron microscopy: S.D. Histology: P.D. Optomotor behavior: D.J. Modeling and data analyses: Z.S., A.D. and M.J. M.J., P.D. and R.C.H. designed the experiments. M.J. wrote the manuscript with all authors contributing in editing. M.J., P.D. and R.C.H. procured funding.

## Acknowledgements

We thank Nigel Atkinson, Allen Shearn, and the Bloomington Stock Centers for reagents. We thank the members of the Juusola lab for discussions and critical readings of the manuscript. This work was supported by the following grants to MJ: Biotechnology and Biological Sciences Research Council (BB/H013849/1, BB/F012071/1 and BB/D001900/1), Engineering and Physical Sciences Research Council (EP/P006094/1), Leverhulme Trust (RPG-2012-567), Jane and Aatos Erkko Foundation, High-End Foreign Expert Grant by Chinese Government (GDT20051100004) and Beijing Normal University (Open Research Fund); grants to RCH: Biotechnology and Biological Sciences Research Council (BB/M007006/1 and BB/J0092531/1), and grants to ZS: 111 Project (NO.B18015), the key project of Shanghai Science & Technology (No.16JC1420402), Shanghai Municipal Science and Technology Major Project (No.2018SHZDZX01) and Zhangjiang Laboratory.

